# Imprecise neural computations as source of human adaptive behavior in volatile environments

**DOI:** 10.1101/799239

**Authors:** Charles Findling, Nicolas Chopin, Etienne Koechlin

## Abstract

Everyday life features uncertain and ever-changing situations. In such environments, optimal adaptive behavior requires higher-order inferential capabilities to grasp the volatility of external contingencies. These capabilities however involve complex and rapidly intractable computations, so that we poorly understand how humans develop efficient adaptive behaviors in such environments. Here we demonstrate this counterintuitive result: simple, low-level inferential processes involving imprecise computations conforming to the psychophysical Weber Law actually lead to near-optimal adaptive behavior, regardless of the environment volatility. Using volatile experimental settings, we further show that such imprecise, low-level inferential processes accounted for observed human adaptive performances, unlike optimal adaptive models involving higher-order inferential capabilities, their biologically more plausible, algorithmic approximations and non-inferential adaptive models like reinforcement learning. Thus, minimal inferential capabilities may have evolved along with imprecise neural computations as contributing to near-optimal adaptive behavior in real-life environments, while leading humans to make suboptimal choices in canonical decision-making tasks.

## Introduction

Everyday life features uncertain situations. In such environments, optimal adaptive behavior requires making probabilistic inferences about external contingencies and especially, about stimulus-action-outcome contingencies. Empirical studies provide ample evidence that such inferential processes operate in the brain and guide human adaptive behavior^1,2,3,4,5,6,7,8,9,10^. As everyday life also features ever-changing situations, optimal adaptive behavior further requires making higher-order probabilistic inferences about the environment *volatility*, i.e. the probability that contingencies change over time. Empirical studies show that consistently, humans adjust their behavior with respect to the environment volatility and in accordance with the computation of such higher-order inferences^1,11,12,13,14^. However, these inferences are complex and may rapidly yield intractable computations^13,15^. This computational complexity problem casts severe doubt upon the hypothesis that such higher-order inferential processes operate in the brain^13^. Thus, we poorly understand how humans exhibit adaptive behavior close to optimal adaptive processes involving inferential computations viewed as biologically implausible.

To clarify this issue, we leveraged the three following facts: (1) in such higher-order inferential processes, inferring a larger environment volatility yields past observations to less strongly influence posterior beliefs about external contingencies^1^; (2) neural computations in inferring external contingencies are highly imprecise, which attenuates the influence of past observations upon posterior beliefs^16^; (3) computational imprecisions further scale with the magnitude of changes in internal representations (the so-called Weber’s Law in psychophysics^17,18,19^). We then hypothesized that imprecise computations in inferring external contingencies dispense with developing higher-order inferences about the environment volatility to optimize adaptive behavior in real-life environments^20^. The rationale is that consistent with the Weber’s law, computational imprecisions in inferring external contingencies are larger in high than low volatile environments. This effect yields past observations in more volatile environments to less strongly influence posterior beliefs, similar to higher-order inferential processes tracking volatility.

To test this hypothesis, we first developed the optimal adaptive processes corresponding to uncertain, stable or changing, closed or open-ended environments. Using advanced machine learning methods, we investigated their performances in these environments. We then demonstrated that the minimal inferential model confined to first-order inferences about external contingencies without tracking volatility exhibits near-optimal adaptive performances in all these environments, provided that computational imprecisions conforming to the Weber’s law corrupt the inferential process. As a result, this *Weber-imprecision* model outperforms no-inferential adaptive processes like reinforcement learning. Secondly, we recorded the performance of human participants in such environments and observed that the Weber-imprecision model accounts for human performances significantly better than (1) the optimal adaptive models, (2) their standard, biologically plausible algorithmic approximations, and (3) reinforcement learning. We then conclude that imprecise neural computations may have been preserved throughout the evolution of minimal, computationally frugal inferential processes as efficiently contributing to near-optimal adaptive behavior in real-life environments, while leading to suboptimal choices in canonical decision-making tasks.

## Results

### Adaptive behavior paradigm

We considered an adaptive agent that responded to successively presented stimuli. In every trial, one among *N* distinct stimuli was randomly drawn and the agent responded by selecting one among *M* actions. The agent then received a positive or negative feedback. The agent thus searched for the correct responses to stimuli by trial and error. However, feedbacks were stochastic and the combination of correct responses to stimuli changed episodically. More precisely, the environment episodically switched across distinct stimulus-response combinations: every combination specified one distinctive response to each stimulus (the correct response) that led to positive feedbacks with unknown, constant probability *η* > 0.5, while the other responses led to positive feedbacks with probability 1 − *η* < 0.5. The maximal number *K* of potential combinations (or latent states) was thus equal to *M*! /(*M*−*N*)!. The current correct combination changed between two successive trials with probability *τ.* Probability *τ* is named the *volatility* and is typically small.

We investigated a range of prototypical environments by first varying number *K* of combinations (**Methods**). *Closed* environments were modeled using *K*=2 combinations corresponding to the repetition of one stimulus (*N*=1) and two available actions (*M*=2), *i.e.* a two-armed bandit with potential reversals between correct actions. *Open-ended* environments were modeled using *K*=24 combinations corresponding to three stimuli (*N*=3) and four available actions (*M*=4), so that uncertainty also bears upon identifying current correct combination: whenever the current combination changed, the new combination *k* was drawn from the set of potential combinations with unknown probability *γ*^*k*^. Secondly, we independently varied the temporal structure of volatility *τ. Stable* environments were modeled by setting volatility to zero (*τ* =0), so that the correct combination remained unchanged across trials. To make the task non-trivial, we considered various stable environments by manipulating feedback sparsity: stable environments delivered feedbacks in 100%, 50%, 20%, 10%, 5% or 2% of trials. *Changing* environments were modeled by setting volatility to a non-zero, either low or large constant, thereby characterizing slow and fast changing environments. Finally, *unstable* environments were modeled by assuming volatility *τ* to follow a bounded, Gaussian random walk (0.03 < *τ* < 0.2), so that the environment volatility smoothly and stochastically varied across trials (**Supplementary Fig. 1**). Note that stable, changing and unstable environments are nested: stable environments are a special case of changing environments, which in turn are a special case of unstable environments.

### Optimal adaptive models

For each prototypical environment described above, we developed the optimal adaptive agent, which inferential process corresponds to the environment generative process. In unstable environments, first, the optimal agent comprises *three* hierarchically-organized levels of inferences^1^ bearing upon: (1) the volatility change rate; (2) the successive values *τ*_*t*_ of the environment volatility; and (3) the successive occurrences of correct combinations *z*_*t*_ (**Fig. 1A**). Congruent with these environments, the agent assumes volatility *τ*_*t*_ to vary as a bounded, Gaussian random walk with (unknown) variance *v*. This agent thus makes nested probabilistic inferences about volatility change rate *v*, environment volatility *τ*_*t*_ and correct combinations *z*_*t*_ based on the successive occurrences of feedbacks. These inferences further combine with those about feedback noise *η* and when *K*>*2*, about occurrence probabilities *γ*^*k*^ of combinations *k*. This whole inferential process results in successively forming posterior beliefs ***B***(*t*) about the current correct combination in trial t. By marginalizing over these beliefs, the agent then selects the most likely correct action in response to stimuli.

**Figure 1.**
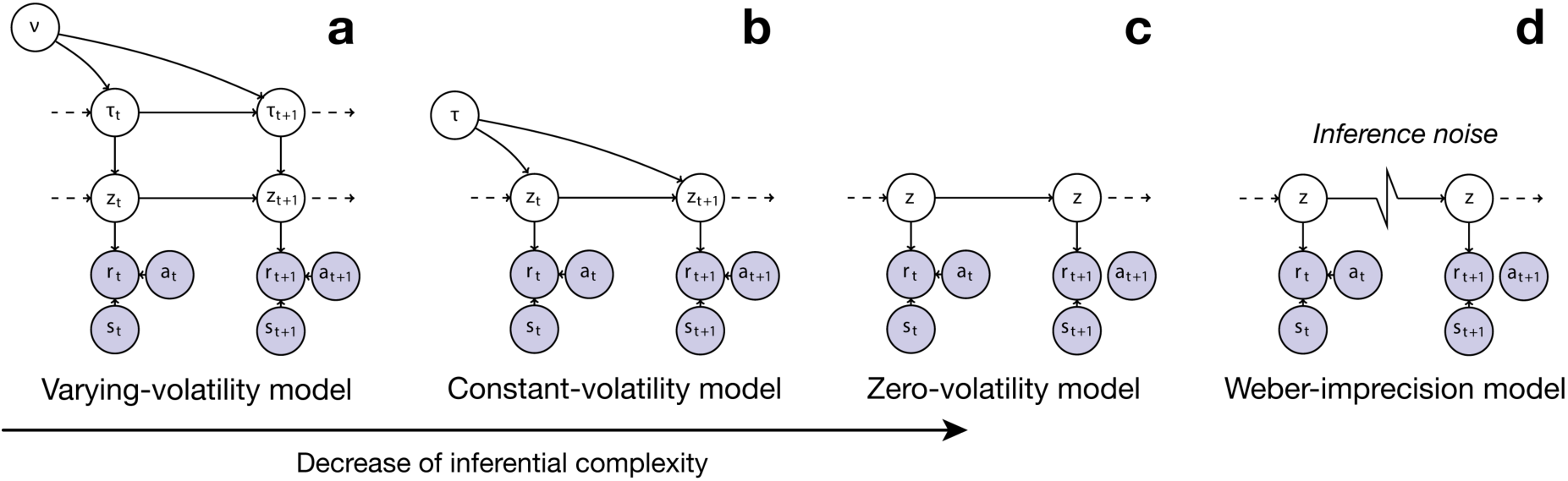
Inferential models of adaptive behavior. **A**, third-order inferential models (referred to as *varying-volatility* models) assuming environment’s latent state *z*_*t*_ in trial *t* changes with probability (volatility) *τ*_*t*_, which in turn is assumed to vary as a Gaussian random walk with constant variance *v.* Inferences bear upon volatility variance *v*, volatility *τ*_t_, and latent state *z*_*t*_ according to external feedbacks *τ*_t_ observed following action a_t_ selected in response to stimuli s_t_. Note that although variance *v* is assumed to be constant, its estimates may change across trials. **B**, Second-order inferential models (referred to as constant-volatility models) assuming environment’s latent state *z*_*t*_ changes with constant probability (volatility) *τ*. Inferences bear only upon volatility *τ* and latent state *z*_*t*_. Again, although volatility *τ* is assumed to be constant, its estimates may change across trials. **C**, first-order inferential models (referred to as zero-volatility models) assuming environment’s latent state *z* remains unchanged across trials. Inferences bear only upon latent state *z*. Again, although latent state *z* is assumed to be constant, its estimates may change across trials. **D**, the Weber-imprecision model is identical to the zero-volatility model except that imprecisions occur in inferential computations. These computational imprecisions reflect neural noise in a manner consistent with the Weber’ law. See text for details. **Supplementary Figs. 3, 4, 5** show the full generative models corresponding to the varying-volatility, constant-volatility and zero-volatility model, respectively.

This agent belongs to the class of inferential models mixing inferences bearing upon both changing latent states *(z*_*t*_ and *τ*_*t*_) and constant, unknown parameter values *(η, γ*_*k*_ and *v*). In such models, computing posterior beliefs is intractable. We therefore emulated this agent using the sequential Monte Carlo method based on particle filtering that has been recently developed in machine learning to solve this issue^21^ **(Methods).** Briefly, posterior beliefs are sequentially sampled through a set of particles that realize iterated Importance Sampling in both the latent-state and parameter space^22,23^ along with an additional, particle-Markov-Chain-Monte-Carlo sampling in the parameter space^24^. We refer to this agent emulation as the *exact varying-volatility* model.

In changing environments, the optimal adaptive agent is identical to the preceding one, except that the agent now comprises only *two* hierarchically-organized inferential levels bearing upon: (1) the value *τ* of the environment volatility, assuming that this value is constant; and as above, (2) the successive occurrences of correct combinations *z*_*t*_) **(Fig. 1B).** This agent belongs to the same class of inferential models as above and is again computationally intractable. We therefore emulated the agent with the same particle filtering method. We refer to this emulation as the *exact constant-volatility* model.

For both models, the particle-filtering method emulates the *exact* inferential process that combines offline both backward and forward inferences: backward inferences enable to revise past beliefs ***B***(*t*<*T)* according to the action outcome observed in current trial *T*, while forward inferences enable to pass these revisions on posterior beliefs ***B***(*T)* in current trial *T*. The exact inferential process therefore lacks biological plausibility and is computationally exorbitant. Consequently, we also investigated the online forward algorithmic approximation of these inferential models: we restricted the particle filtering described above to the online iterated Importance Sampling that estimates posterior beliefs ***B***(*T*) from posterior beliefs ***B****(T-1)* in the preceding trial^22,23^**(Method).** The resulting online particle filtering is plausibly implemented in populations of cortical neurons^25,26,27^. We refer to these approximations as the *forward* varying-/constant-volatility model, respectively.

In stable environments, finally, the optimal adaptive agent is identical to the preceding agents except that it now comprises only *one* inferential level bearing upon external contingencies **(Fig. 1C):** namely the identity of the correct combination *z*, assuming that this combination remains unchanged over trials *(i.e.* the environment volatility is assumed to be zero). The resulting inferential complexity is considerably lower: in contrast to the preceding models, this agent is computable through pure online forward closed-form computations^28^: in every trial t, posterior beliefs ***B***(*T*) about combination z can be directly computed from posterior beliefs ***B***(*t*-1) in trial t-1 according to the action outcome observed in trial *t* **(Methods).** For consistency, however, we emulated the agent using the same forward particle-filtering method as above knowing that in this case, the forward approximation emulates the exact inference process. We refer to this emulation as the *zero-volatility* model.

### The Weber-imprecision model

The zero-volatility model has the critical advantage to rely on computable online forward inferences but evidently, has poor adaptive performances in volatile environments when combinations change: posterior beliefs ***B***(*t*) about the correct combination can only move slowly from the current to the next correct combination. We hypothesized that the substantial amount of computational imprecisions previously identified in human inferential processes^16^ remedies this limitation. Consistent with the psychophysical Weber’s law^17,18^, indeed, computational imprecisions presumably scale with the magnitude of belief updating and consequently, increase whenever combinations change. Such computational imprecisions make posterior beliefs ***B***(*T*) less dependent upon past observations, especially when combinations more frequently change: posterior beliefs ***B****(t*) thus adjust to the environment volatility as if they derive from higher-order inferences about the environment volatility. We hypothesized that with such imprecisions, the zero-volatility model approaches the optimal adaptive performance in volatile environments.

Computational imprecisions presumably derive from the noise in online neural computations. We consequently assumed that in the *(forward)* zero-volatility model, particle-filtering is noisy: every particle coding for one combination in trial *t* may start (mis)coding for another combination in trial *t*+1 with probability *ϵ*_*t*_. Consistent with the Weber’s law, computational imprecisions are further assumed to scale with the distance *d*_*t*_ between posterior beliefs in trial *t* and *t+1.* We therefore assumed noise *ϵ*_*t*_ is a random variable uniformly distributed between 0 and *μ* + *λd*_*t*_ :

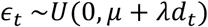

where *μ* ≥ 0 and *λ* ≥ 0 are free parameters quantifying the constant and Weber component of computational imprecisions, respectively **(Methods).** We refer to the zero-volatility model comprising such inferential imprecisions as the *Weber-imprecision* model **(Fig. 1D)**. For *μ* = 0 and *λ* = 0, the Weber-imprecision model simply reduces to the zero-volatility model. **Supplementary Fig. 2** shows that as expected, only when : is nonzero, posterior beliefs ***B***(*T*) are sensitive to the environment volatility as if they derive from higher-order inferences about the environment volatility.

### Alternative models

To assess the functional specificity of computational imprecisions in inferences, we investigated an alternative model whereby computational imprecisions occur in action selection rather than in belief inferences. This model, referred to as the *Noisy-Selection* model, is identical to the zero-volatility model, except that actions are probabilistically selected according to the standard softmax rule with inverse temperature *β* as free parameter. Thus, for *β* ≫ 1, the noisy-selection model becomes identical to the zero-volatility model.

Finally, to assess the role of inferential processes in optimizing adaptive behavior, we also considered the standard, non-inferential adaptive model, namely the reinforcement learning model *(RL)* combining the Rescorla & Wagner’s rule and softmax rule with learning rate *α* and inverse temperature *β* as free parameters, respectively^29,30^ **(Methods).**

### Computer simulation results

We simulated all the models in every environment described above and comprising 1000 trials. We set feedback noise *η* to 90%. In each environment (characterized by number *K*, its volatility structure and feedback sparsity), we simulated every model 50 times and computed its resulting average performance (proportion of correct responses). In all unstable environment simulations, we set volatility variance *v* to 0.0001. In slow and fast changing environments, constant volatility *τ* was set to *τ*_*l*ow_=0.03 and *τ*_high_=0.2, respectively. In every open-ended environment simulation, occurrence probabilities *γ*^*k*^ of combinations were set randomly. Chance level was 50% and 25% in closed *(K=2)* and open-ended *(K=24)* environments, respectively. We first analyzed the best performance that the Weber-imprecision, noisy-selection and RL model could achieve in every environment. For every environment, we thus computed the free parameters that maximize model performances. We then compared these best performances to the corresponding optimal agent performances: namely the performance of the exact zero-, constant- and varying-volatility models in stable, changing and unstable environments, respectively or equivalently, to their forward approximation, which performance was virtually identical to the exact models **(Fig. 2).**

**Figure 2.**
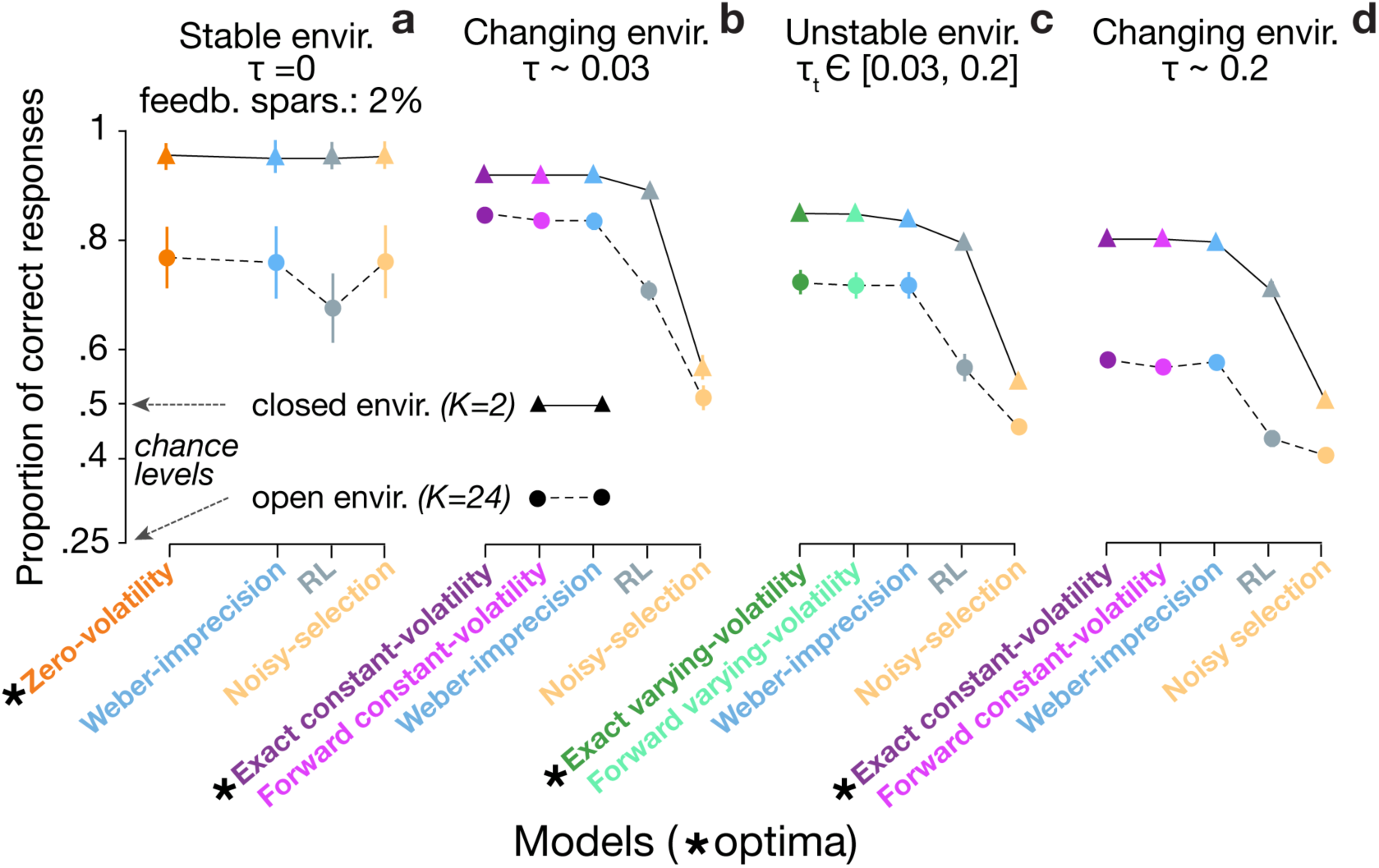
Models’ maximal performances in stable, changing and unstable environments. Maximal proportions of correct responses for the Weber-imprecision, Reinforcement-learning (RL) and Noisy-selection model when stimulated in closed (K=2 latent states, triangle) and open-ended (K=24 latent states, circle) environments. Each model was simulated N=50 times in every environment (error bars are s.e.m. across simulations).* corresponds to the theoretical optimal model for each environment. **A**, stable environments (volatility *τ* = 0) with sparse feedbacks (2% of trials). The theoretical optimal performance corresponds to the zero-volatility model. **B**, slow changing environments (constant volatility *τ* = 0.03). The theoretical optimal performance corresponds to the exact constant-volatility model. **C**, unstable environments (volatility *τ*_t_ follows a bounded Gaussian random walk in the range [0.03; 0.2]). **D**, fast changing environments (constant volatility *τ* = 0.2). The theoretical optimal performance corresponds to the exact constant-volatility model. Note that in every environment, the forward approximation of the exact optimal model as well as the Weber-imprecision model virtually achieved the optimal performance. By contrast, the RL performed decently in closed environments but poorly in open-ended environments, while the Noisy-selection model (zero-volatility model comprising a softmax bearing upon action selection) performed poorly in both closed and open-ended environments.

In stable environments, as expected, both the Weber-Imprecision and Noisy-Selection model performed optimally, regardless of feedback sparsity and number *K* of latent states (optimal performances when feedback sparsity was 2%: 96% and 77% for *K*=2 and *K*=24, respectively; s.e.m.<2%) **(Fig. 2A).** Both models indeed comprise the zero-volatility model as special cases. By contrast, model RL reached the optimal performance only when *K*=2. When *K*=24 and feedback sparsity was equal to 2%, model RL exhibited a 10% loss of performance (s.e.m.=3%), reflecting the lack of inferential processes about latent states.

In changing environments as hypothesized, only the Weber-imprecision model performed quasi-optimally **(Fig. 2B,D).** For *K*=2, the optimal performance was 92.1% and 80.4% (s.e.m.<0.2%) in low (*τ*_low_∼0.03) and high volatile (*τ*_high_∼0.2) environments respectively, while the Weber-imprecision model performance was 92.0% and 79.7% respectively (s.e.m.<0.2%). For *K*=24, the optimal performance was 85% and 58% in low and high volatile environments, respectively (s.e.m.<0.3%), while the Weber-imprecision model performance was 83% and 57%, respectively (s.e.m.<0.4%). Model RL performed moderately for *K*=2 (89% and 71% for *τ*_low_ and *τ*_high_, respectively, s.e.m.<0.3%) but poorly for *K*=24 (71% and 44% for *τ*_low_ and *τ*_high_, respectively, s.e.m.<0.6%). Finally, the Noisy-selection model performed even more poorly for both *K*=2 and *K*=24, regardless of the environment volatility (<57% and 51% for *K*=2 and *K*=24, respectively, s.e.m.<0.7%) **(Fig. 2B,D).**

In unstable environments, the results also confirmed our prediction **(Fig. 2C).** Only the Weber-imprecision model still performed quasi-optimally. For *K*=2 and *K*=24, the optimal performance was 85.1% (s.e.m.=0.3%) and 72.3% (s.e.m.=0.7%) respectively, while the Weber-Imprecision model achieved a performance of 84.0% (s.e.m.=0.3%) and 71.6% (s.e.m.=0.8%), respectively. Again, model RL performed moderately for *K*=2 (80%) but poorly for *K*=24 (57%) (both s.e.m.<0.8%) while in both cases, the Noisy-selection model exhibited a dramatically poor performance (54% and 46%, respectively, s.e.m.<0.5%).

Unlike the noisy-selection and RL model, thus, the Weber-imprecision model virtually reached the optimal performance in a variety of stable and volatile environments. This optimal performance however was reached with noise parameters (*μ, λ*) properly adjusted to each environment. This adjustment deviates from the idea that as reflecting neural computational imprecisions, these parameters are precisely not adjustable, especially through higher-order inferential processes. To address this issue, we next investigated the *versatility* of model performances across all the environments considered above with no parameter adjustments. For that purpose, we first computed for every pair of noise parameters (*μ, λ*) the performance loss that in each environment, the Weber-imprecision model exhibited relative to the optimal performance. For each pair (*μ, λ*), we then computed the maximal performance loss (denoted maxloss(*μ, λ*)) over all the environments. **Fig. 3A** shows that consistent with our prediction, maxloss(*μ, λ*) remained below 10% when constant noise component 8 remained close to zero (<0.04) and Weber noise component : ranged between ∼0.8 and ∼1.8. Maxloss(*μ, λ*) was minimal and equal to 9.2% loss (s.e.m.=0.16%) when *μ*\* = 0.01 and *λ*\* = 1.4. In every environment, accordingly, the Weber-imprecision model with parameters (*μ*\*, *λ*\*) exhibited a performance loss relative to the optimal performance that never exceeded 9.2% (s.e.m.=0.16%). This upper bound value, which we refer to as the *minimaxloss*, defines the model versatility over environments.

**Figure 3.**
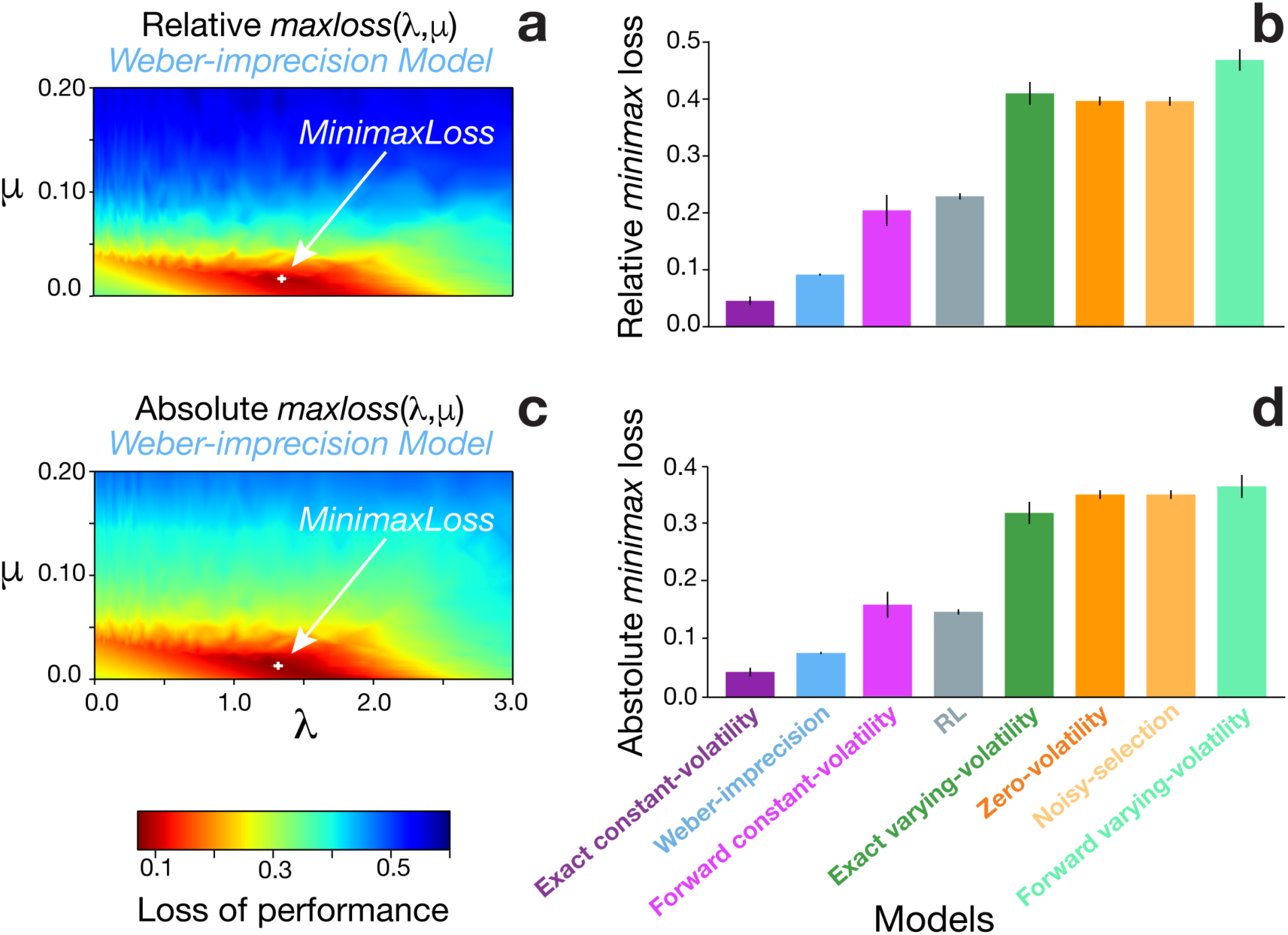
Models’ versatility across environments. **A**, maximal losses over all investigated environments (stable, slow and fast changing, unstable, either closed or open-ended) of the Weber-imprecision model relative to theoretical optimal performances according to noise free parameters *μ* and *λ* (constant and Weber noise components, respectively). These losses are normalized in each environment with respect to the related theoretical optimal performance. The arrow indicates the *minimaxloss* that corresponds to the parameter values minimizing these maximal losses. **B**, minimaxlosses for all the investigated models. **C**, same as **A**, except that absolute rather than relative losses are considered: namely, losses in every environment are not normalized with respect to the related theoretical optimal performance. **D**, same as **B** except that minimax losses are computed from absolute rather than relative losses. Each model was simulated N=50 times in each environment (error bars are s.e.m. across simulations). The lower the minimaxloss, the more the model is versatile across the environments. Note that the exact constant-volatility model is the most versatile, but is biologically implausible from a computational viewpoint. Among the biologically plausible models (Weber-imprecision, RL, zero-volatility, noisy-selection, forward constant-/varying-volatility model), the Weber-imprecision is the most versatile and approaches the versatility of the exact constant-volatility model.

Using the same minimax approach, we next investigated the RL and Noisy-selection models versatility. We found that these models were dramatically less versatile than the Weber-imprecision model. The RL model’s minimaxloss was equal to 22.7% (s.e.m.=0.53%) and corresponded to learning rate *α\** = 0.83 and inverse temperature *β\** = 12.3, *i.e.* to almost a Win-Stay, Loose-Switch policy **(Fig. 3B).** For the Noisy-selection model, the minimaxloss was equal to 39.6% (s.e.m.=0.78%) and corresponded to inverse temperature *β*\* → +∞ *i.e.* to the zero-volatility model **(Fig. 3B).** Thus, the Weber-imprecision model versatility could neither be achieved through RL processes nor noisy action selection but specifically resulted from imprecise probabilistic inferences over external contingencies.

We next compared the Weber-imprecision model versatility to both the constant- and varying-volatility model versatility. The latter models are presumably highly versatile, as the environments are nested: stable environments correspond to changing environments with a constant volatility equal to zero, while changing environments correspond to unstable environments with a zero-variance Gaussian random walk. These models comprise no free parameters, so that the minimaxloss simply reduces to the maxloss these models exhibited across the environments. We found that only the *exact constant-* volatility model was more versatile than the Weber-imprecision model **(Fig. 3B).** While the latter exhibited a 9.5% minimaxloss (s.e.m.=0.16%) as reported above, the minimaxloss for the exact constant-volatility model was 4.5% (s.e.m.=0.76%). Its forward approximation (*i.e.* the forward constant-volatility model) however was as poorly versatile as the RL model (minimaxloss=20.3%, s.e.m.=2.72%), while both the exact and forward varying-volatility performed even more poorly (minimaxloss>40%, s.e.m.<2%). All these higher-order inference models exhibited the largest performance loss in environments with sparse feedbacks indicating that: (1) when feedbacks are sparse, third-order inferences about volatility change rate *v* are highly inefficient and even deleterious, as the number of volatility trajectories between two distant informative trials are potentially infinite; (2) assuming a constant environment volatility overcomes this problem but requires backward processes to bring together distant feedbacks for properly inferring the environment volatility. Finally, it is worth noting that the Weber-imprecision model exhibited its worst performance relative to the optimal one (reaching its maximal relative loss=9.5%) in the environment where the constant-volatility model is precisely the optimal agent (changing environment), volatility is both large (high volatility *τ*_high_∼0.2) and represents the only source of uncertainty regarding correct combinations (closed environment *K*=2). Among the biologically plausible models, the Weber-imprecision model thus appeared as the most versatile adaptive model with a performance loss relative to the theoretical adaptive optima that remained below 10%. We obtained the same results when considering the absolute rather than relative performance losses **(Fig. 3C,D).**

### Human adaptive performances in closed environments

The results above lend theoretical support to the hypothesis that the Weber-imprecision model describes neural computations guiding human adaptive behavior in the variety of real-life environments. We investigated this hypothesis by first testing 22 participants in a behavioral protocol implementing the unstable, closed environment described above *(i.e. K*=2, two-armed bandits with reversals) **(Fig. 4A)**. One participant was excluded because her/his overall performance was lower than two standard deviations from the mean performance. The protocol exactly corresponded to this environment except that: volatility *τ*_*t*_ varied as a step-wise function rather Gaussian random walk, taking its values in the set [.01, .02, .03, .05, .08, .15] every 120 trials in an order counterbalanced across participants **(Fig. 4B);** Feedback noise *η* was set to 80% rather than 90%. We made these changes to make the protocol similar to that used in previous studies^1^. Model computer simulations showed that in this protocol, the exact varying- and constant-volatility model performances were 86.9% and 86.3%, respectively (s.e.m.<0.3%). Their forward approximation reached virtually the same performances (86.6% and 86.0%, respectively; s.e.m.<0.3%). Participants’ performances was significantly lower (79.8%, s.e.m.=0.9%), although as previously shown^1^, their behavior was sensitive to the environment volatility: immediately after nonregular trials (negative feedbacks after correct responses and vice-versa along with reversals), participants’ responses like models’ responses changed more frequently in high than low volatile episodes (median split; two-tailed paired *T*-test; participants and models: all *Ts*(20)>2.9, all *Ps*<0.01; **Fig. 4C**).

**Figure 4.**
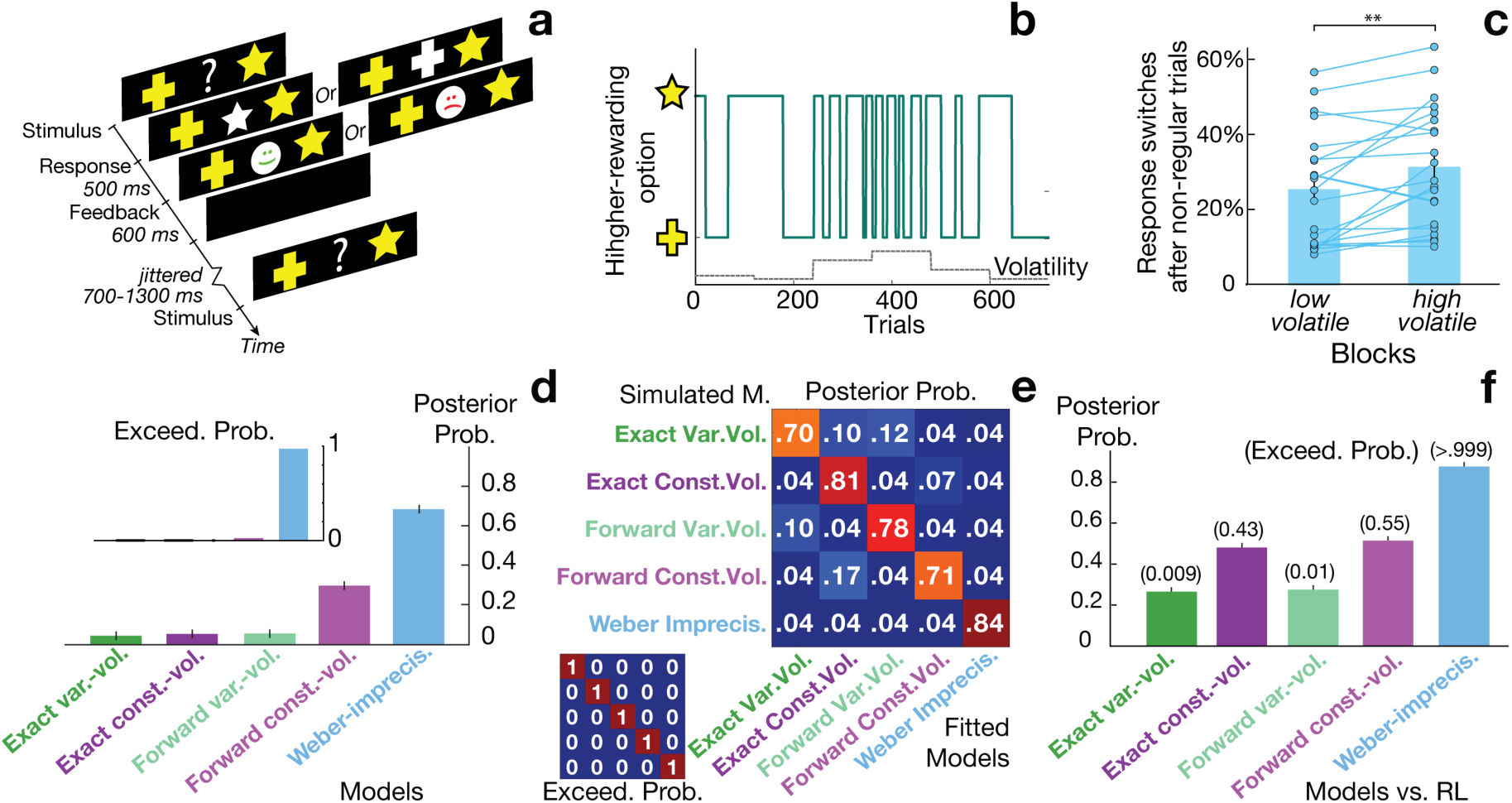
Model fits to human performances in closed, unstable environments. **A**, trial structure in the two-armed bandit task with stochastic feedbacks (closed environment). In every trial, participants chose one between two visually presented symbols by pressing a response button and received a binary, positive or negative feedback. One symbol led to positive feedbacks with probability 80% (correct response), the other led to negative feedback with probability 80%. **B**, temporal structure of the two-armed bandit task. The more frequently rewarded symbol changed episodically (reversals). The volatility (probability of reversals) varied along the experimental session as a step-wise function comprising six distinct levels (unstable environment) which occurrence order was counterbalanced across subjects. Each level comprised 120 trials. The panel shows one example of experimental sessions with this temporal structure. **C.** Proportion of participants’ response switches after non-regular trials (i.e. comprising trials when the computer delivered the low-probability (P=20%) feedback and trials featuring reversals), in low vs. high volatile blocks (median split). Participants switched more often in high than low volatile blocks, indicating that they adapted their behavior according to the environment volatility (**p<0.01, two-tailed paired T-test). **D**, estimated model posterior probabilities with respect to human data and related exceedance probabilities (Inset) for inferential models (varying- and constant-volatility models, their forward approximation and the Weber-imprecision model). The Weber-imprecision model best accounted for human performances decisively (*P*_*exceed*_ *=* 0.973). **E**, confusion matrices from the model recovery procedure assessing the protocol ability to dissociate the models. The large matrix shows estimated model posterior probabilities with respect to synthetic behavioral data generated by each model with free parameters fitted to each participant (*N* = 21). The small matrix shows the related exceedance probabilities and reveals that each model decisively explains its own behavior better than the other models. **F**, estimated posterior probabilities of each model relative to the reinforcement learning model (RL) with respect to human data. Related exceedance probabilities are shown in parenthesis. Only the Weber-imprecision model decisively accounts for human data better than RL. In all these analyses, action selection in every model is assumed to follow a softmax probabilistic function (free parameter: inverse temperature *β*). See **Supplementary Tables 1,2,3** for model fitted parameters.

To fit participants’ choice data, we inserted in every model a softmax rule for action selection (free parameter: inverse temperature *β*). Note that as a result, the Weber-imprecision model included the Noisy-selection model as a special case with noise parameters set to zero. To compare model fits, we derived model posterior probabilities (MPPs) given the data from computing model *marginal* likelihoods over parameter spaces using advanced Monte-Carlo procedures **(Methods).** MPPs based on marginal likelihoods are the optimal Bayesian quantification for selecting models, balancing model degree of freedom and adequacy to data. Among all inferential models, as expected, the optimal but biological implausible models (exact varying-/constant-volatility models) showed the lowest MPPs **(Fig. 4D).** The forward varying-volatility model fitted hardly better. In contrast, the forward constant-volatility and Weber-imprecision MPPs were about 5 and 12 times as large as this baseline, respectively. Thus, the Weber-imprecision model showed the largest MPPs, which further led to reject the other inferential models (Exceedance probability: *P*_*exceed*_ = 0.973). Consistent with the simulation results, the best-fitting Weber-imprecision model (*i.e.* with free parameters maximizing model likelihoods) relied on the dominant contribution of the Weber noise component (*λ*_*fit*_⟨*d*_*t*_⟩ *=* 0.25; *μ*_*fit*_ = 0.05, s.e.m.=0.01; *λ*_*fit*_ = 0.8, s.e.m.=0.1) and as participants, switched its response immediately after non-regular trials more frequently in high than low volatile episodes (median split; two-tailed paired *T*-test: *T*(20)=2.6, *P*=0.019).

To confirm these findings, we verified using a model recovery procedure **(Methods)**^31^ that the behavioral protocol associated with our fitting method properly discriminated the models. According to MPPs, indeed, every model fitted its own performance simulated from its best-fitting free parameters better than the other models and led to reject them (all *Ps* _*exceed*_ >0.99, **Fig. 4E**). Moreover, we found that only the Weber-imprecision model explained participants’ data better than the RL model (*P*_*exceed*_>0.999), while the other models failed (all *Ps*_*exceed*_≤0.55, **Fig. 4F**). Consistently, the best-fitting Weber-imprecision model fully captured the time-course of participants’ performances following reversals, while the other best-fitting models failed **(Fig. 5).**

**Figure 5.**
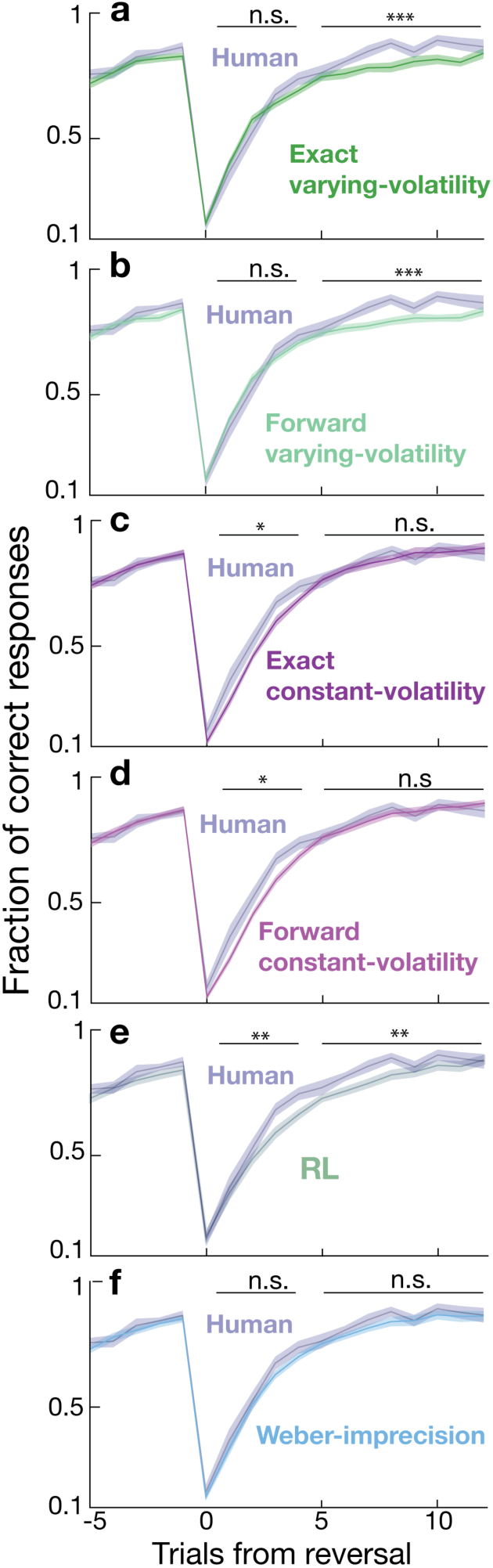
Human and model adaptive behavior following contingency reversals. Fraction of correct responses in trials preceding and following reversals in the two-armed bandit task described in **Fig. 4** (closed, volatile, unstable environment). Each graph shows human performances (same data in all graphs) vs. fitted model performances averaged across participants (shaded areas are s.e.m. across participants: N=21). Trial #0 corresponds to the trial when unbeknownst to participants and models, reversals occurred. Human performances showed fast early adaptive responses to reversals (trials 1 to 3) followed by a slow, late adaptation to new environment contingencies (trials 5 to 15) leading to a performance plateau. **A**, human vs. exact varying-volatility model performances. Late adaptations (averaged over trials 5 to 15) significantly differed between the two (two-tailed paired T-test, *T*(20) = 3.9, *p* < 0.001). B, human vs. forward varying-volatility model performances. Again, late adaptations significantly differed between the two (paired T-test, *T*(20) = 4.7, *p* < 0.001). **C**, human vs. exact constant-volatility model performances. Early adaptive responses significantly differed between the two (two-tailed paired T-test, *T*(20) = 2.39, *p =* 0.027). **D**, human vs. forward constant-volatility model performances. Again, early adaptive responses significantly differed between the two (two-tailed paired T-test, *T*(20) = 2.65, *p* = 0.015). **E**, human vs. reinforcement learning (RL) performances. Both early adaptive responses and late adaptations differed the two (two-tailed paired T-tests, both *Ts*(20) > 2.91, *p* < 0.008). **F**, human vs Weber-imprecision model performances. The Weber-imprecision model exhibited early adaptive responses and late adaptations similar to participants (two-tailed paired T-tests, both *Ts*(20) < 1.12, *p* > 0.12). Thus, constant-volatility models responded to reversals slower than participants, varying-volatility models exhibited late adaptations to new environment contingencies slower than participants, RL responded to reversals and adapted to new contingencies slower than participants. Only the Weber-imprecision model exhibited adaptive performances similar to participants. * *p*< 0.05, \*\**p*< 0.01, *** *p*< 0.001, n.s.= not significant.

### Human adaptive performances in open-ended environments

We next tested whether the Weber-imprecision model accounts for human adaptive behavior in open-ended environments. We collected the adaptive performances from 62 participants tested in two previously published studies^5,7^. These participants were tested in a behavioral protocol identical to the slow changing, open-ended environment described above (constant volatility *τ*_low_=0.03, K=24) **(Fig. 6A).** Unbeknownst to participants, every participant performed two distinct instantiations of the environment: in one instantiation, every combination occurred once, thus forming 24 episodes associated with the 24 distinct combinations *(no-recurrence* condition); in the other instantiation, the 24 episodes corresponded to distinct combinations pseudo-randomly drawn from a subset of only three combinations *(recurrence* condition). Model computer simulations over these two instantiations showed that in this protocol, the optimal performance, *i.e.* the exact constant-volatility model performance, was equal to 86.9% (s.e.m.=0.4%). The exact varying-volatility model reached virtually the same performance (86.7%, s.e.m.=0.4%). Their forward approximation also achieved virtually the same performance (86.1% and 85.4%, respectively; s.e.m.<0.4%). Participants exhibited a significantly lower performance (74.2%, s.e.m.=0.7%).

**Figure 6.**
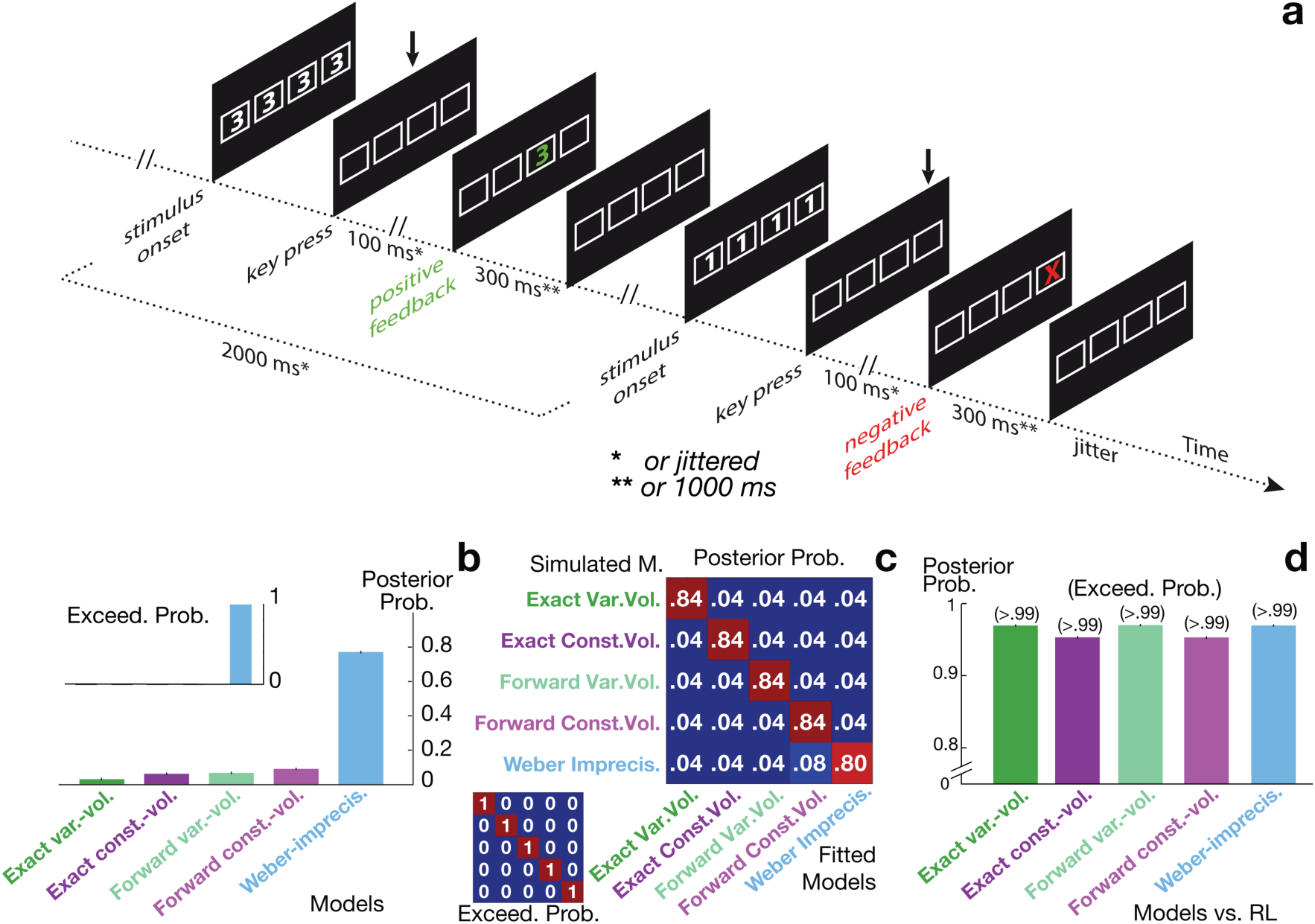
Model fits to human performances in open-ended, changing environments. **A**, trial structure in the open-ended task, modeling slow changing environments with stochastic, positive or negative feedbacks. Visual stimuli were pseudo-randomly drawn from a set of three Arabic numbers, e.g. (1,3,5). Subjects responded by pressing one among four possible response keys. Shortly after subjects’ responses, stimuli were removed and a positive or negative feedback was presented. Each digit was associated with a unique correct response leading to positive feedbacks with probability 90%. The other responses led to negative feedbacks with probability 90%. Digit-response combinations changed episodically and unpredictably with constant volatility *t* =0.03. Two successive combinations were orthogonal, i.e. all stimulus-response pairs were distinct. Adapted from^5,7^ * and ** indicates slight timing differences in trial structures between^5^ and^7^ due to specific neuroimaging constraints in the latter study. **B**, estimated model posterior probabilities with respect to human data from^5,7^ and related exceedance probabilities (Inset) for inferential models (varying- and constant-volatility models, their forward approximation and the Weber-imprecision model). The Weber-imprecision model best accounted for human performances decisively *(P*_*exceed*_ > 0.99). **C**, confusion matrices from the model recovery procedure assessing the protocol ability to dissociate the models. The large matrix shows estimated model posterior probabilities with respect to synthetic behavioral data generated by each model with free parameters fitted to each participant (*N* = 62). The small matrix shows the related exceedance probabilities and reveals that each model decisively explains its own behavior better than the other models. **D**, estimated posterior probabilities of each model relative to the reinforcement learning model (RL) with respect to human data. Related exceedance probabilities are shown in parenthesis. All inferential models decisively account for human data better than RL. In all these analyses, action selection in every model is assumed to follow a softmax probabilistic function (free parameter: inverse temperature *β*). See **Supplementary Tables 1,2,3** for model fitted parameters.

To fit participants’ choice data, we again inserted in every model a softmax rule for action selection. In these open-ended environments, computing model marginal likelihoods was practically intractable. To derive MPPs, we therefore used model Bayesian Information Criteria (BIC) as standard estimates of marginal likelihoods, with model free parameters maximizing model likelihoods **(Methods).** We first found that every inferential model (varying-/constant-volatility models; Weber-imprecision model with the Noisy-selection model as a special case) explained participants’ data better than model RL (all *Ps _exceed_* >0.99) **(Fig. 6D).** Unlike participants and inferential models, indeed, model RL is unable to exploit the re-occurrence of combinations in the recurrence condition^5^. Yet, the same model recovery procedure as above confirmed that the protocol associated with our fitting method properly discriminated the inferential models: according to MPPs, indeed, every model fitted its own performance simulated from its best-fitting free parameters better than the other models and led to reject them (all *Ps* _*exceed*_ >0.99, **Fig. 6C)**. Critically, the Weber-imprecision model was again found to best explain participants’ data **(Fig. 6B)**: the Weber-imprecision MPP was at least 8 times as large as both the exact and forward varying-/constant-volatility MPPs, which led to reject these volatility-based inference models (*P*_*exceed*_ >0.99). Consistent with the preceding results, the best-fitting Weber-imprecision model again relied on the dominant contribution of the Weber noise component (*λ*_*fit*_⟨*d*_*t*_⟩ = 0.39; *μ*_*fit*_ = 0.05, s.e.m.=0.002; *λ*_*fit*_ = 1.2, s.e.m.=0.06).

## Discussion

Optimal adaptive behavior in uncertain and variable environments is based on making first-order probabilistic inferences about stimulus-action-outcome contingencies along with higher-order probabilistic inferences about the environment volatility. We observed that as previously reported^1,11,12,13,14,32,33,34^ human adaptive performances in some volatile environments more likely reflected such higher-order inference processes than reinforcement learning, while in other volatile environments, the converse was observed. In both cases, however, we found evidence that first-order inferential processes enduring computational imprecisions accounted for human adaptive performances significantly better than both higher-order inference processes and reinforcement learning. The same results were found even when considering biologically plausible approximations of higher-order inference processes that exclude offline backward inferences. Moreover, the results show that these computational imprecisions corrupting the formation of posterior beliefs conformed to the general Weber’s law by scaling with the magnitude of belief updating^17,18,19^. These findings provide an empirical support to the hypothesis —named the *Weber-imprecision* model— that imprecise computations in first-order inferences about stimulus-action-outcome contingencies guide human adaptive behavior in uncertain and variable environments.

Our computer simulation results provide a principled support to the Weber-imprecision model. First of all, the simulations show this counter-intuitive result: in a variety of environments (closed, open-ended and with zero, constant or unstable volatility), the Weber-imprecision model can virtually reach the *adaptive optimum* in every environment, *i.e.* the adaptive performance derived from the inferential process matching the environment generative process and that outperforms reinforcement learning. This indicates that the Weber-imprecision model captures key adaptive properties of optimal processes which in previous studies, may have been (mis)interpreted as evidence supporting the presence of higher-order inferential adaptive processes in humans.

Second, our computer simulations show that among the biologically plausible models involving no backward inferences, the Weber-imprecision model exhibits adaptive performances emerging as the most versatile/robust to *structural uncertainty*^12^ (also named *Knightian* or *Keynesian* uncertainty in economics^35,36^). As frequent in real-life, structural uncertainty reflects the lack of knowledge about the generative structure of the environment. We considered here environments with three distinct temporal structures: stable (zero-volatility), changing (constant volatility) and unstable (random-walk volatility) environments. As the unstable generative structure encompasses the changing one, which in turn encompasses the stable one, in principle, the third-order inference process corresponding to the adaptive optimum in unstable environments optimally unravels this structural uncertainty. However, when action feedbacks were sparse as often the case in everyday situations, we found this adaptive process to exhibit performances lower than reinforcement learning (maximal performance losses relative to adaptive optima >23%; see Results). This finding confirms that as previously advocated^12,15^, resolving temporal structural uncertainty through higher-order inferences bearing upon the actual, highdimensional space of possible temporal structures is actually ineffective and even deleterious in view of the informative poverty of environment feedbacks.

In contrast, the second-order inference process corresponding to the adaptive optimum in changing environments approached the adaptive optimum in *every* environment, even when feedbacks were sparse (maximal performance losses relative to adaptive optima <5%). This adaptive process assumes a constant volatility and in principle, optimally unravels the structural uncertainty only across changing and stable environments. Nevertheless, as posterior beliefs about the volatility value are constantly updated, the process efficiently adapts to unstable environments. With sparse feedbacks, moreover, assuming a constant volatility rules out the considerable variety of volatility trajectories compatible with successive and temporally distant feedbacks but inferentially inextricable. Within all the environments investigated here, this second-order inference process thus represents the best compromise between the dimensionality and adequacy of the inferential space. However, its biologically plausible approximation excluding offline backward inferences exhibited adaptive performances not better than reinforcement learning, when feedbacks were sparse (maximal performance losses relative to adaptive optima > 20%, see Results). This result indicates that second- and higher-order inferential processes require offline backward inferences for bridging across no-feedback trials to properly infer the environment volatility and for exhibiting adaptive performances surpassing reinforcement learning.

Although the Weber-imprecision model only relies on (forward) first-order inferences corresponding to the adaptive optimum in stable environments, we found this model to resolve structural uncertainty close to the full second-order inference process even when feedbacks were sparse (maximal performance losses relative to adaptive optima <10%), thereby outperforming both the forward approximation of the latter and reinforcement learning. The Weber-imprecision model indeed assumes external contingencies to remain constant across trials which in any volatile environments, remains the default and “irrefutable” hypothesis in absence of environment feedbacks or when environment feedbacks remain consistent with posterior beliefs. Moreover, when external contingencies change and generate feedbacks inconsistent with these beliefs, the Weber’s computational imprecisions enable posterior beliefs to rapidly detach from prior ones and converge towards new contingencies. Thus, the computer simulations reveal that among the biologically plausible processes, the Weber-imprecision model is the most versatile and robust to the temporal structural uncertainty: this model approaches the best adaptive performances, which the offline, biologically implausible second-order inferential process achieve.

The Weber-imprecision model, however, is not an algorithmic approximation of higherorder inference processes. Indeed, the model performances maximally differed in environments when either the former or the latter corresponded to the adaptive optimum. Conceptually, the Weber-imprecision model optimizes short-term predictability by rapidly inferring locally accurate but globally inaccurate simplified world models (stable contingencies), while computational imprecisions enable to rapidly switch across such simplified world models. By contrast, higher-order inferential processes optimize longterm predictability by slowly inferring more general, accurate but complex world models (volatile contingencies), detrimental to short-term effectiveness.

The Weber-imprecision model relies on online first-order inferences about external contingencies that can be expressed in closed form to compute posterior beliefs in trial *t+1* directly from those in trial t. We nonetheless chose to implement these inferences through an online particle-filter based on forward, iterated Importance-Sampling^22,23^ for four reasons: (1) this particle-filter realizes the exact first-order inference; (2) it constitutes the forward inference component in the particle-filtering method we used to emulate higher-order inferential processes^21^, which allowed us to properly compare the inferential models; (3) it is neurally plausible, as previous studies have shown this particle-filter may reflect neural computations^25,26,27^; (4) In online particle-filtering, computational imprecisions are naturally modeled as sampling noise that scales with the extend of resampling that occurs between successive trials. As this particle-filter emulates the exact first-order inference, this implementation choice has no impacts on the results: the same results will be obtained with other exact implementations of firstorder inferences, like evidence accumulation models and their neural implementations reviews in^37,38^, along with computational imprecisions consistent with the Weber’s law. The present results thus remain agnostic about possible neural implementations underlying the Weber-imprecision model.

In any cases, our Weber-imprecision model assumes that as previously proposed^19^, computational imprecisions consistent with the Weber’s law stem from intrinsic neural noise and corrupt the inferential process. We indeed modeled computational imprecisions as a sampling noise that arises *randomly and uniformly* over a range scaling with the magnitude of belief updating between successive trials. An algorithmic-equivalent model assuming no computational imprecisions would *compute* and monitor this magnitude online to deterministically scale the sampling noise accordingly. To decipher between these two interpretations, we again compared models’ posterior probabilities using the same procedure as described in Results given the human data we collected in the closed and open-ended environments. We found significant evidence favoring the computational imprecision interpretation in both environments (both *P*s_exceed_ >0.99). This confirms the Weber-imprecision model that the variability of human performances relative to the first-order inference process reflects imprecise neural computations rather than additional monitoring processes. In canonical decision-making tasks featuring stable environments, such imprecise neural computations act as a nuisance. For instance, Drugowitsch et al.^16^ showed that these imprecisions accounts for 2/3 of human suboptimal choices in standard tasks requiring participants to infer from a series of ambiguous cues which among several latent states is the more likely. The present results however show these imprecisions endow humans with powerful adaptive abilities in volatile environments, which approach the optimal but biologically implausible adaptive performance.

In all the inference models considered here, we assumed first-order inferences about external contingencies to bear upon the whole set of latent states. This is certainly an unrealistic assumption in open-ended environments as those investigated here (*K*=24 potential combinations). Using the same behavioral data set, we previously showed that participants actually make first-order inferences bearing only upon a small subset of three/four latent states referred to as the *inferential buffer*^5,7^. These behavioral data along with associated neuroimaging data thus support a model, named model *Probe*, describing how this inferential buffer through these first-order inferences is updated with new latent states and drives adaptive behavior, *i.e.* the learning and selection of combinations guiding behavior^5,7^. However, this model as originally proposed treats the environment volatility as a constant free parameter modulating first-order inferences as deriving from higher-order inferences. The present results therefore predict that replacing this free parameter with computational imprecisions consistent with the Weber’s law should provide an even better account of observed human performances. To test this prediction, we used the same procedure as described in Results and given human data, we compared model posterior probabilities associated with the original Probe model and the revised one assuming such computational imprecisions occur within the inferential buffer **(Methods).** We found model posterior probabilities to significantly favor the revised model Probe (*P*_*exceed*_ > 0.99). This result shows that the Weber-imprecision model accounts for human adaptive performances, without requiring first-order inferences to bear upon all the latent states of the environment.

Neuroimaging results show that in unstable environments, the dorsomedial prefrontal cortex exhibits activations correlating with the volatility derived from third-order inference processes realizing the adaptive optimum^1^. At first sight, this finding provides some support that higher-order inference processes guide human adaptive behavior. As shown here however, the Weber-imprecision model is able to virtually achieve this adaptive optimum. This indicates that the related computational imprecisions influence first-order inferences about external contingencies in a way similar to the inferred volatility. Accordingly, dorsomedial prefrontal activations correlating with the inferred volatility might actually reflect the discrepancy between the resulting imprecise beliefs truly encoded in this region to guide participants’ choices and the exact beliefs derived from third-order inferential models. This hypothesis makes a testable prediction for future research: as mentioned above indeed, imprecise neural computations in the Weber-imprecision model further exhibit stochastic fluctuations independent of the volatility derived from higher-order inference processes.

In conclusion, previous results have shown that imprecise neural computations in human inferential processes about external contingencies largely contribute to human choice suboptimality in canonical decision-making tasks^16^. The present results show that these imprecisions in first-order inferences actually dispense humans from developing higher-order inferences about the environment volatility to reach near-optimal adaptive behavior in volatile environments. Thus, imprecise neural computations may have been preserved throughout the evolution of computationally frugal inferential processes as efficiently contributing to near-optimal adaptive behavior in real-life environments.

## Methods

### Adaptive behavior paradigm

We considered an adaptive agent responding to successively presented stimuli. In every trial, one among *N* distinct stimuli was randomly drawn uniformly and the agent responded by selecting one among *M* actions. The agent then received a positive or negative feedback. The agent thus searched for the correct responses to stimuli by trial and error. However, feedbacks were stochastic and the combination of correct responses to stimuli changed episodically. More precisely, the environment episodically switched across distinct “latent states” defining the current correct combination: every latent state specified one distinctive response to each stimulus (the correct response) that led to positive feedbacks with unknown, constant probability *η* > 0.5, while the other responses led to positive feedbacks with probability 1 – *η* < 0.5. The maximal number *K* of potential latent states (or correct combinations) was thus equal to *M*! /(*M* – *N*)!. Latent states changed between two successive trials with probability *τ* named volatility. We simulated the following environments:

1. *Closed and stable environments.* These environments correspond to *K*=2 and volatility *τ* =0, namely a two-armed bandit with no reversals: a unique stimulus (*N*=1) was repeated over trials and one between two available actions (*M*=2) had to be selected in every trial. To make such environments non-trivial and investigate the role of feedback sparsity on adaptive behavior, we simulated eight closed and stable environments each delivering feedbacks in 100%, 50%, 20%, 10%, 5% or 2% of trials. Feedback noise *η* was set to 90%.
2. *Closed and changing environments:* these environments were identical to the preceding closed and stable environments except that volatility *τ* was non-zero and constant. This corresponds a two-armed bandit with reversals between two latent states. We simulated slow and fast changing environments corresponding to constant volatility *τ* = 0.03 and *τ* = 0.2, respectively. As such environments are non-trivial, feedbacks were delivered in every trial (feedback sparsity was set to zero).
3. *Closed and unstable environments.* These environments were identical to the preceding closed and changing environments except that volatility *τ*_*t*_ varied across trials as a bounded Gaussian random walk within the range [0.03,0.2] and with variance *v* = 0.0001: in every trial *t*, volatility *τ*_*t*_ is drawn from a normal distribution centered on volatility *τ*_*t*-1_ with variance *v* (and whenever the variable was drawn lower or larger than 0.03 and 0.2, respectively, volatility *τ*_*t*_ was set to 0.03 and 0.2, respectively).
4. *Open-ended and stable environments.* These environments were identical to the closed and stable environments described above except that number *K* of potential latent states was set to *K*=24: In every trial, one among three stimuli (*N*=3) was drawn randomly and uniformly and one among four available actions (*M*=4) had to be selected, which leads to *K*=24 potential combinations. The correct combination remained unchanged across trials. Again, we simulated eight open-ended and stable environments each delivering feedbacks in 100%, 50%, 20%, 10%, 5% or 2% of trials.
5. *Open-ended and changing environments.* These environments were identical to the preceding open-ended and stable environments except that volatility *τ* was non-zero and constant. As above, we considered slow and fast changing environments corresponding to constant volatility *τ* = 0.03 and *τ* = 0.2, respectively. As such environments are non-trivial, feedbacks were delivered in every trial (feedback sparsity set to zero). When latent states changed, the new latent state *k* is drawn according to a multinomial distribution with probabilities *γ*^1^, …, *γ*^*k*^, …,*γ*^*K*^ (excluding the preceding latent state). Probabilities were initially drawn randomly and uniformly according to a flat Dirichlet distribution (in order that Σ_*i*=1,…*K*_ *γ*^*i*^ = 1).
6. *Open-ended and unstable environments.* These environments were identical to the preceding open-ended and changing environments except that volatility *τ*_*t*_ varied across trials as a bounded Gaussian random walk within the range [0.03,0.2] and with variance *v* = 0.0001, as described above for closed, unstable environments.

### Computational models

#### Exact varying-volatility model

The exact varying-volatility model corresponds to the optimal Bayesian adaptive process with a generative model exactly matching the generative process of *unstable* environments described above with *K* equal to either 2 (closed environments) or 24 (open environments) (see **Fig. 1A** and **Supplementary Fig. 3**). This model is therefore the theoretical adaptive optimum in these environments. More precisely, the model assumes that:

i. volatility *τ*_*t*_ varies as a bounded Gaussian random walk within the range [0,0.5] with uninformative priors about constant variance *v*:

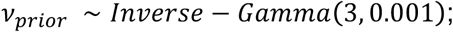
ii. the current latent state changes between trials *t-1* and *t* with probability *τ*_*t*_;
iii. when the current latent state changes between trial *t-1* and *t*, new latent state *z*_*t*_ is drawn from the set of potential latent states {1,…,*K*} according to a multinomial distribution with probabilities ***γ*** = {*γ*^1^, …, *γ*^*K*^}:

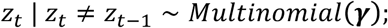

with Σ_*k*=1,…*K*_ *γ*^*k*^ = 1 and excluding the preceding state;
iv. priors about probabilities ***γ*** are uninformative and follow a flat Dirichlet distribution:

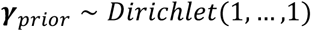
v. Given current latent state *z*_*t*_, stimulus *s*_*t*_ and chosen action *a*_*t*_, feedbacks *r*_*t*_ are delivered according to a Bernoulli distribution with parameters *η* or 1 – *η*:

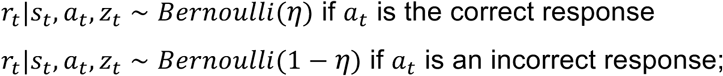
vi. priors about feedback noise *η* > 0.5 are uninformative and follow a flat Beta distribution:

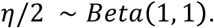

By marginalizing over posterior beliefs about latent states *z*_*t*_, the model then selects in every trial the action that most likely leads to the positive feedback in response to the stimulus *(i.e.* the “correct” action): namely, actions are selected according to an “argmax” policy. In this model, however, computing posterior beliefs about latent states *z*_*t*_, *τ*_*t*_ and latent parameters *v*, ***γ***, *η* is an intractable problem. We addressed this issue by using a sequential Monte-Carlo (SMC) algorithm recently developed in machine learning to solve this class of inferential models comprising both latent states and parameters^21^. The algorithm is based on particle filtering methods and converges to the exact solution when the number of sampling particles increases to infinity^21^. The algorithm comprises two intermixed SMC procedures: (1) a particle filter^22^ implementing iterated Important Sampling in the space of latent states (*z*_*t*_, *τ*_*t*_); (2) an iterated Importance Sampling combined with a particle Markov Chain Monte Carlo method^24,39^ in the parameters’ space *v*, ***γ***, *η*. We implemented the algorithm using a total number of 10^6^ particles corresponding to 1000 samples in the parameters’ space, each associated with 1000 particles in the space of latent spaces. We verified that this number allows approaching the asymptotic convergence: we implemented the algorithm using 4 × 10^6^ particles and obtained virtually identical posterior beliefs.

#### Exact constant-volatility model

The exact constant-volatility model corresponds to the optimal Bayesian adaptive process with a generative model exactly matching the generative process of *changing* environments described above with *K* equal to either 2 (closed environments) or 24 (open environments) (see **Fig. 1B** and **Supplementary Fig.4**). This model is therefore the theoretical adaptive optimum in these environments. More precisely, the exact constant-volatility model is identical to the exact varying-volatility model except that volatility *τ*_*t*_ is now assumed to be a constant (*τ*_*t*_ = *τ* for all *t*) in the range [0,0.5]. Priors about volatility *τ* were uniform over [0,0.5] and followed a flat Beta distribution:

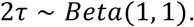

The exact constant-volatility model was implemented using the same SMC algorithm as the exact varying-volatility model described above.

#### Forward varying- and constant-volatility models

As explained above, both the exact varying- and constant-volatility models perform exact inference through particle filtering methods (when the number of particles is large enough). However, the algorithm is both computationally very costly and biologically implausible because it constantly requires offline backward passes to re-sample and revise past beliefs about latent parameters according to all past observations: beliefs’ re-sampling and revisions are performed in a backward fashion through the Particle Markov Chain Monte Carlo (P-MCMC) procedure to prevent the sampling of beliefs from degenerating in local minima and to more accurately sample posterior beliefs. To overcome this limitation, we derived an online, biologically plausible, computationally frugal approximate algorithm from the exact SMC algorithm described above. This forward approximation replaces the offline, backward P-MCMC procedure with the following standard, online particle resampling of posterior beliefs: resampling occurs with respect to the current empirical estimates of parameters’ distributions given parameter priors. For the varying-volatility model, accordingly, the resampling of posterior beliefs about latent parameters *v*, ***γ***, *η* occurs in trial *t* through *Inverse-Gamma, Dirichlet* and *Beta* distributions, respectively, and which hyper-parameters are estimated from the particle filter in trial *t* (**Supplementary Methods**). For the constant-volatility model, similarly, the resampling of posterior beliefs about latent parameters *τ*, ***γ***, *η* occurs in trial *t* through *Beta, Dirichlet* and *Beta* distributions, respectively, and which hyper-parameters are again estimated from the particle filter in trial *t.* In both cases, the resulting online particle filtering is biologically plausible and neural implementations have been previously proposed^25,26,27^. We referred to these approximations as the *forward* varying-/constant-volatility model, respectively. To make these forward models as computationally frugal as possible, we implemented them with the minimal number of particles that allows approaching the asymptotic performance in the most complex environment (*i.e.* unstable environments): namely 200 × 200 = 40000 particles (see **Supplementary Fig. 6**). Overall, the results show that these forward models provide accurate approximations of exact models, provided that the environment and model generative process are identical (**Figs. 2** and **3**).

#### Zero-volatility and Noisy-selection model

The zero-volatility model corresponds to the optimal Bayesian adaptive process with a generative model exactly matching the generative process of *stable* environments described above with *K* equal to either 2 (closed environments) or 24 (open environments) (see **Fig. 1C** and **Supplementary Fig. 5**). This model is therefore the theoretical adaptive optimum in these environments. More precisely, the zero-volatility model is identical to the exact constant-volatility model except that volatility *τ* is now assumed to be zero. Model priors thus reduce only to combination probabilities ***γ*** and feedback noise *η* as described above. Posterior beliefs in this model are computable in closed form^28^. For consistency, however, we implemented this model through the same forward particle filtering method as that used for the forward varying-/constant-volatility models, knowing that in this case, the forward approximation emulates the exact inference process. The Noisy-selection model is the variant of the zero-volatility model assuming that actions are selected according to a softmax rather than argmax policy. The Noisy-selection model thus has the softmax inverse temperature *β* as free parameter and comprises the zero-volatility model as a special case (*β* ≫1).

#### Weber-imprecision model

The Weber-imprecision model is another variant of the zero-volatility model assuming that imprecisions stemming from online neural noise occur in computing posterior beliefs about latent states. Neural noise is naturally introduced in the zero-volatility model by implementing a noisy particle filter: every particle coding for one latent state (*i.e.* combinations) in trial *t* may start (mis)coding for another latent state in trial *t*+1 with probability *ϵ*_*t*_. This newly (mis)encoded state is randomly drawn according to the actual distribution of particles over latent states, which reflects probabilities ***γ***. Consistent with the Weber’s law, computational imprecisions are further assumed to scale with the distance *d*_*t*_ between posterior beliefs in trial *t* and *t+1.* Probabilistic noise *ϵ*_*t*_ is thus itself a random variable uniformly distributed between 0 and *μ* + *λd*_*t*_:

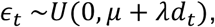

where *μ* and *λ* are two free parameters quantifying the constant and Weber component of computational imprecisions, respectively. **Supplementary methods** show how distance *d*_*t*_ derives from particle filtering. Note that for *μ* = 0 and *λ* = 0, the Weber-imprecision model simply reduces to the zero-volatility model.

#### Reinforcement learning model

The reinforcement learning model combines the Rescorla-Wagner learning rule^29^ (learning rate *α* as free parameter) with a softmax policy for action selection (inverse temperature *β* as free parameter). In response to successive stimuli *s*_*t*_, accordingly, the softmax policy selects the action *a*_*t*_ according to action values Q(*s*_*t*_, *a*), which are updated following feedbacks *r*_*t*_ with respect to the Rescorla-Wagner learning rule:

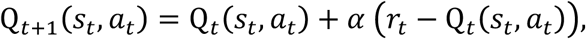

while Q_*t*_ (*s, a*) remain unchanged for *s ≠ s*_*t*_ or *a ≠ a*_*t*_.

### Experimental procedures: closed environments

#### Participants

We recruited 22 healthy participants (including 11 females and 11 males, aged 18-35 years), who volunteered to participate to the study. Participants provided written informed consent. The present study was approved by the French National Ethics Committee (CPP, Inserm protocol #C15-98). Participants were paid for their participation. One participant was excluded because her/is overall performance was lower than two standard deviations from the mean performance.

#### Behavioral protocol

Each participant completed the two-armed bandit task described in **Fig. 4A,B**. Stimuli were presented on a computer screen and participants responded using two response buttons. The task included 720 trials broken down into 6 blocks of 120 trials corresponding to six distinct levels of external volatility. Stimuli remained on the screen until participants responded (self-paced task). 500ms later, the positive or negative feedback (green happy vs. red angry smiley faces) was displayed during 600ms. Inter-Trial Interval was jittered between 700 and 1300ms. Participants had short breaks and were informed that breaks had no influences on task contingencies and had to keep in mind their beliefs about these contingencies during breaks. Final pay-offs increased with the number of positive feedbacks participants received.

### Experimental procedures: open-ended environments

We analyzed the behavioral performances of participants to two previously published studies that administered the same behavioral protocol with and without functional magnetic resonance imaging^5,7^. The behavioral protocol corresponded to the open-ended, slow changing environment described in the main text with constant volatility *τ* = 0.03 (see **Fig. 6A**). The experimental procedures are described in detail in these articles.

#### Collins et al. ‘s study^5^

22 healthy volunteers (13 females; age range: 18-35 years old) participated to the study. No participants were excluded. Stimuli were visually presented Arabic numbers. Participants responded to each stimulus by pressing one of four response keys. When key presses occurred no later than 1500ms after stimulus onsets, stimuli disappeared 100ms after key presses and subjects received audiovisual feedbacks (duration 300ms). Feedbacks were positive or negative. A positive feedback consisted of an ascending sound and the apparition of the associated stimulus in a box representing the pressed key at the bottom of the screen. Negative feedbacks consisted of a descending sound, only. Otherwise, stimuli were removed and no feedbacks were delivered. Stimulus onset asynchrony was 2000ms. Associations between actual stimuli, response fingers and feedbacks were orthorgonalized and counterbalanced across subjects. Participants were instructed that feedbacks could be uncertain and variable and that payoffs increased with the total number of received positive feedbacks. No additional instructions were provided to participants.

In every trial, a “correct” response was associated with each stimulus (three possible stimuli) and led to positive feedbacks with probability 90%. All other responses led to negative feedbacks with probability 90%. Distinct stimuli were associated with distinct correct responses. Digit-response combinations remained unchanged over series of successive trials uniformly ranging from 36 to 54 trials and corresponding to an external volatility *τ* = 0.03. Two successive combinations were orthogonal, *i.e.* all digit-response pairs differed between successive combinations.

The experiment included two behavioral sessions administered in two separate days. In each session, combinations changed 24 times. In the *no-recurrence* session, combinations never re-occurred. In the *recurrence* session, only three distinct combinations occurred in a pseudo-randomized order and in equal proportion. These combinations were fully orthogonal. Stimuli were pseudo-randomly chosen from the set {1,3,5} (one session) or {2,4,6} (the other session). Stimuli along with combination and session order were counterbalanced across subjects. Volatility of external contingencies was identical in the no-recurrence and recurrence sessions (*τ* = 0.03%).

#### Donoso et al. ‘s study^7^

40 healthy, right-handed volunteers (20 females; age range: 18-26 years old) participated to the study (fMRI study). No participants were excluded. The experimental protocol was identical to that in Collins et al.’s study described above, except the following differences. Positive feedbacks corresponded to stimuli presented in green in the box representing the chosen button. Negative feedbacks corresponded to a red cross shown in the box representing the chosen button. When no answer was made, red lines were shown in the four boxes. Pseudo-randomized time jittering was used between responses and feedbacks (from 400 to 3900ms) as well as between feedback offsets and trial onsets (from 100 to 3600ms). Feedbacks duration was 1000ms. (see **Fig. 6A**).

### Model fitting and comparison

In both closed and open-ended environments, we replaced the “argmax” with a softmax decision policy (free parameter: inverse temperature *β*) in all models to fit models’ to participants’ performances. Thus, the exact/forward varying-/constant-volatility models included a unique free parameter: namely, the softmax inverse temperature *β*. The Weber-imprecision model included three free parameters: the softmax inverse temperature *β* along with noise parameters *μ, λ*. As a result, the Weber-imprecision model comprised the zero-volatility and Noisy-Selection model as special cases. Finally, the Reinforcement Learning model included two free parameters: the softmax inverse temperature *β* along with learning rate *α*. Model fits to human data were compared according to Model Posterior Probabilities (MPPs), *i.e.* the model marginal likelihoods (given the data) with uniform priors over models. MPPs based on marginal likelihoods are the optimal Bayesian quantification for comparing models, balancing model degree of freedom (or complexity) and adequacy to data. Model marginal likelihoods are obtained by marginalizing model likelihoods over the whole free parameter space. This marginalization was practically tractable for closed environments and was carried out through standard Importance Sampling and Quasi Monte Carlo methods^40^. For open-ended environments, however, it was practically intractable (especially for the Weber-imprecision model comprising three free parameters) and we used the standard approximation of model marginal likelihoods, *i.e.* the Bayesian Information Criterion (BIC). Computing BICs requires estimating *maximal* model likelihoods over the free parameter space, which was carrying out through standard Bayesian optimization methods^41^.

In any case, computing model likelihoods given free parameter values is required before marginalizing or maximizing these likelihoods. As the inferential models are particle filters, computing the likelihoods for these models is not trivial and is explained below.

#### Exact varying- and constant- volatility model

For these models, computing model likelihoods given human data is straightforward: For each model, indeed, the particle filter emulates the corresponding *exact* inferential process so that model posterior beliefs and consequently, the model softmax probability to select the actual human choices are unrelated to particle filtering. Given inverse temperature *β*, thus, each model directly provides its likelihood given human data.

#### Forward varying- and constant- volatility model

For each of these models, the particle filter approximates and may diverge from the corresponding exact inferential process. As a result, model posterior beliefs and consequently, the model softmax probability to select the actual human choices depend upon the actual realization of the particle filter. Given inverse temperature *β*, computing the model likelihood given human data then requires marginalizing over the realizations of particle filtering. We solved this marginalization problem by noting that as a particle filter, each forward model actually defines a hidden Markov chain generating actions. We therefore computed the marginalization using established sequential Monte Carlo methods for hidden Markov models^23^. Note that this marginalization procedure again optimally balances the adequacy to data and the additional degree of freedom resulting from the various possible realizations of particle filtering.

#### Weber-imprecision model

For this model, the particle filter reflects the *exact* zero-volatility inferential process but the presence of computational imprecisions make the Weber-imprecision model to diverge from the exact process. As a result, model posterior beliefs and consequently, the model softmax probability to select the observed human choices again depend upon the actual realization of the noisy particle filter implementing the Weber-imprecision model. We solved this issue exactly as described above for the forward varying-/constant-volatility model. Given inverse temperature *β* and noise components *μ, λ*, we thus derived the model likelihood given human data. Note that again the derivation is a marginalization procedure and therefore optimally balances the adequacy to data and the additional degree of freedom resulting from the various possible realizations of the noisy particle filter.

### Model recovery procedure

We implemented a *model recovery* procedure to assess the validity of our model fitting and selection procedure and the ability of our experimental protocols to discriminate the models^31^. The recovery procedure consists in generating synthetic data by simulating every model of interest, then applying our model fitting and selection procedure (see above) to these data. This procedure should lead to select the simulated model. For each model, we generated 21 sets (62 sets) of synthetic data in the closed (open-ended, respectively) environment, each corresponding to the free parameters fitted to one participant. The results show the recovery procedure was conclusive for both the closed and open-ended environment, thereby validating our model fitting and selection procedure along with the protocol ability to discriminate the models.

### Model Probe

Model Probe is an online algorithm approximating optimal adaptive processes *(i.e.* Dirichlet processes mixture) in environments featuring a potentially infinite number of latent states^5,7^. Model Probe accounts for human adaptive behavior in open-ended, changing environments as those investigated here^5,7^. Model Probe assumes first-order inferences bear only upon a small subset of three/four latent states referred to as the *inferential buffer*^5,7^ and describes how this inferential buffer is updated with new latent states through these first-order inferences and drives adaptive behavior. The model however treats external volatility as a free parameter modulating first-order inferences as deriving from higher-order inferences, which allows expressing the inferential process in closed-form. To overcome this limitation, we investigated a new model Probe by replacing this free parameter with Weber imprecisions bearing upon first-order inferential computations within the inferential buffer. As Model Probe is expressed in closed-form, these imprecisions were modeled as follow:

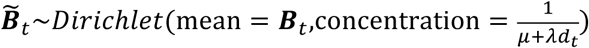

where ***B***_*t*_ are posterior beliefs about latent states inferred according to the original model Probe with volatility parameter set to zero 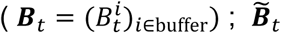 are the corresponding imprecise beliefs resulting from Weber imprecisions. As in the **main text**, 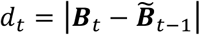 measures the magnitude of belief updating, while *μ* and *λ* are free parameters reflecting the constant and Weber component of neural noise. The Dirichlet distribution appropriately indicates that 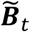 are randomly distributed around mean ***B***_*t*_ with variance scaling with *μ + *λ*d*_*t*_. We fitted and compared the original and new Probe model to human performances recorded in the open-ended environment investigated in **Results**, using the same model fitting and comparison method as for the Weber-imprecision model (see section **model fitting and selection** above). Model posterior probabilities (MPPs) revealed that the new Probe model decisively best accounts for human performances (exceedance probability *P*_*exc*_ > 0.99).

## Supplementary Information

**Supplementary Figure 1:**
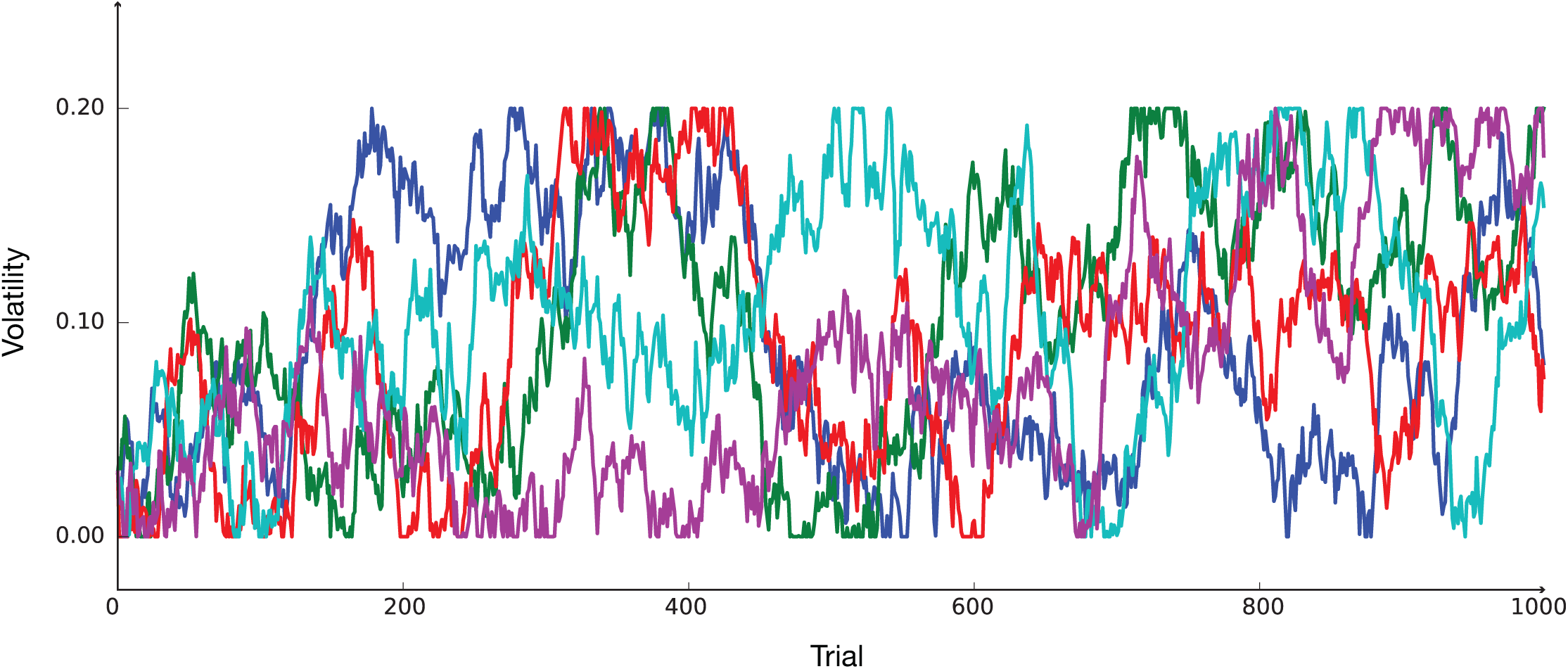
Examples of five volatility trajectories in unstable environments. Environment volatility follows a bounded random walk between 0.03 and 0.2 with variance 0.0001 (see Methods).

**Supplementary Figure 2:**
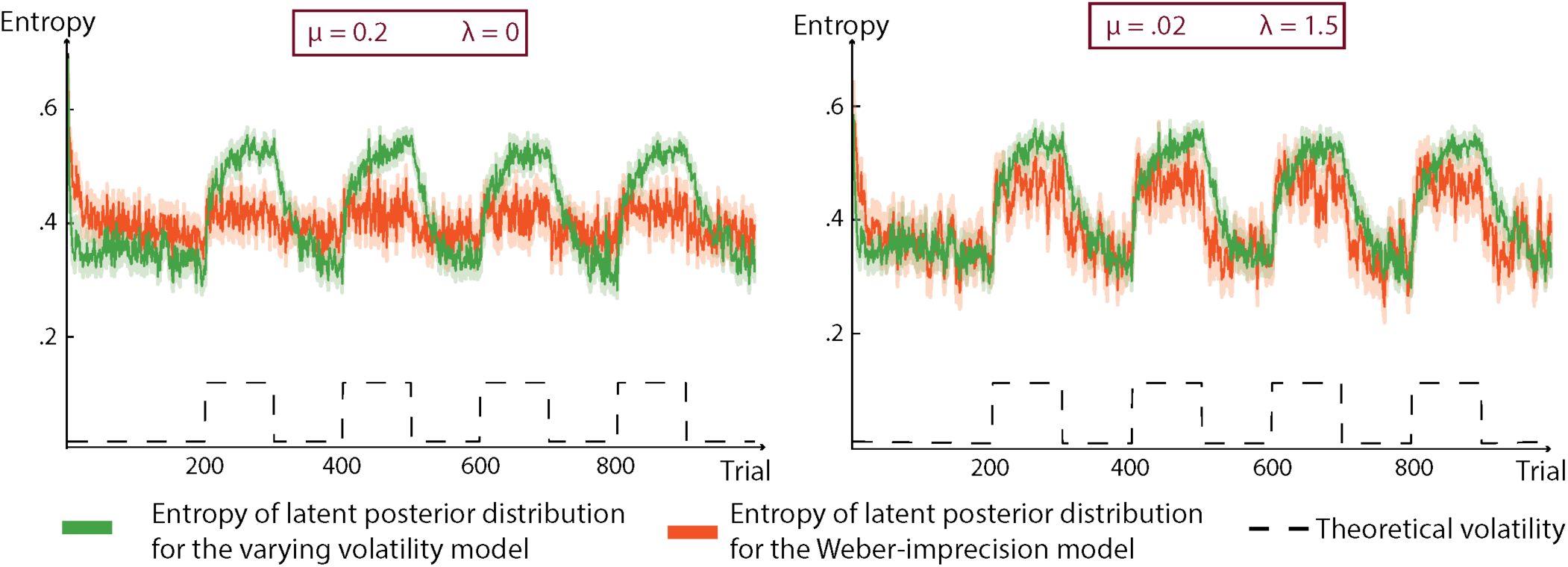
Relation between the Weber noise component λ in the Weber-imprecision model and external volatility. The figure shows the entropy of posterior beliefs about current combinations (latent state posteriors) for the exact varying-volatility and Weber-imprecision model. Each model is simulated *N* = 50 times in a closed environment (K=2, two-armed bandit), which alternates between high and low volatility periods. Left, simulations when Weber component *λ* is set to 0 and constant component *µ* is large (*µ* = 0.2). Right, simulations when Weber component A is large (*λ* = 1.5) and constant component *µ* is low (*µ* = 0.02). Note that the entropy of posterior beliefs are similar between the Weber-imprecision and varying-volatility model only when the Weber component is large enough.

**Supplementary Figure 3:**
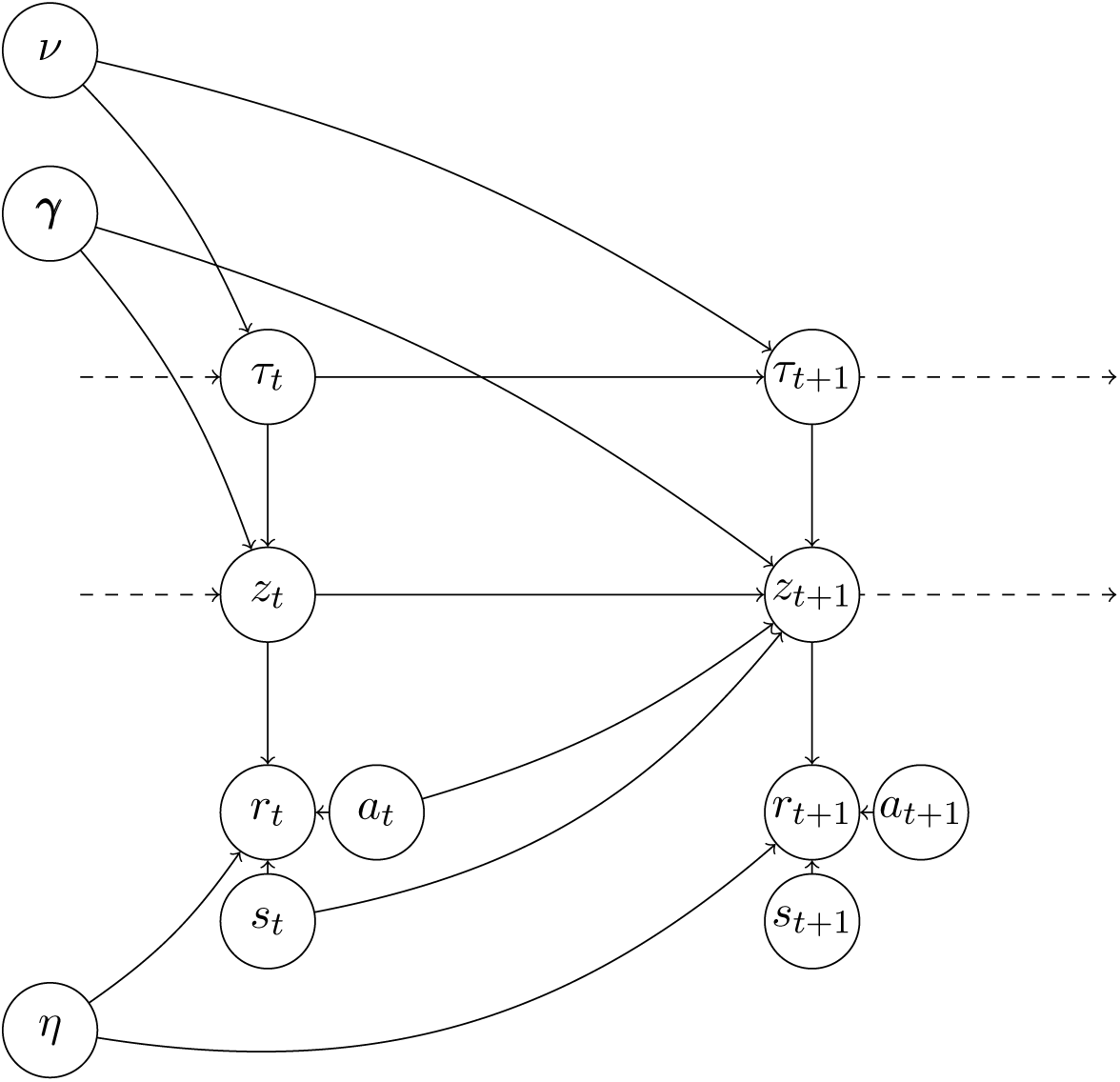
Full generative model of varying-volatility models. This model is exactly the process generating *unstable* environments. This generative model assumes that volatility *τ*_*t*_ follows a bounded random walk with constant variance *ν. z*_*t*_ represents the current correct combination. *γ* represents the probabilities of combination occurrence whenever the correct combination changes. *η* represents feedback noise. In every trial, observables are stimuli *s*_*t*_, actions *a*_*t*_ and binary feedback *r*_*t*_. See Methods for details.

**Supplementary Figure 4:**
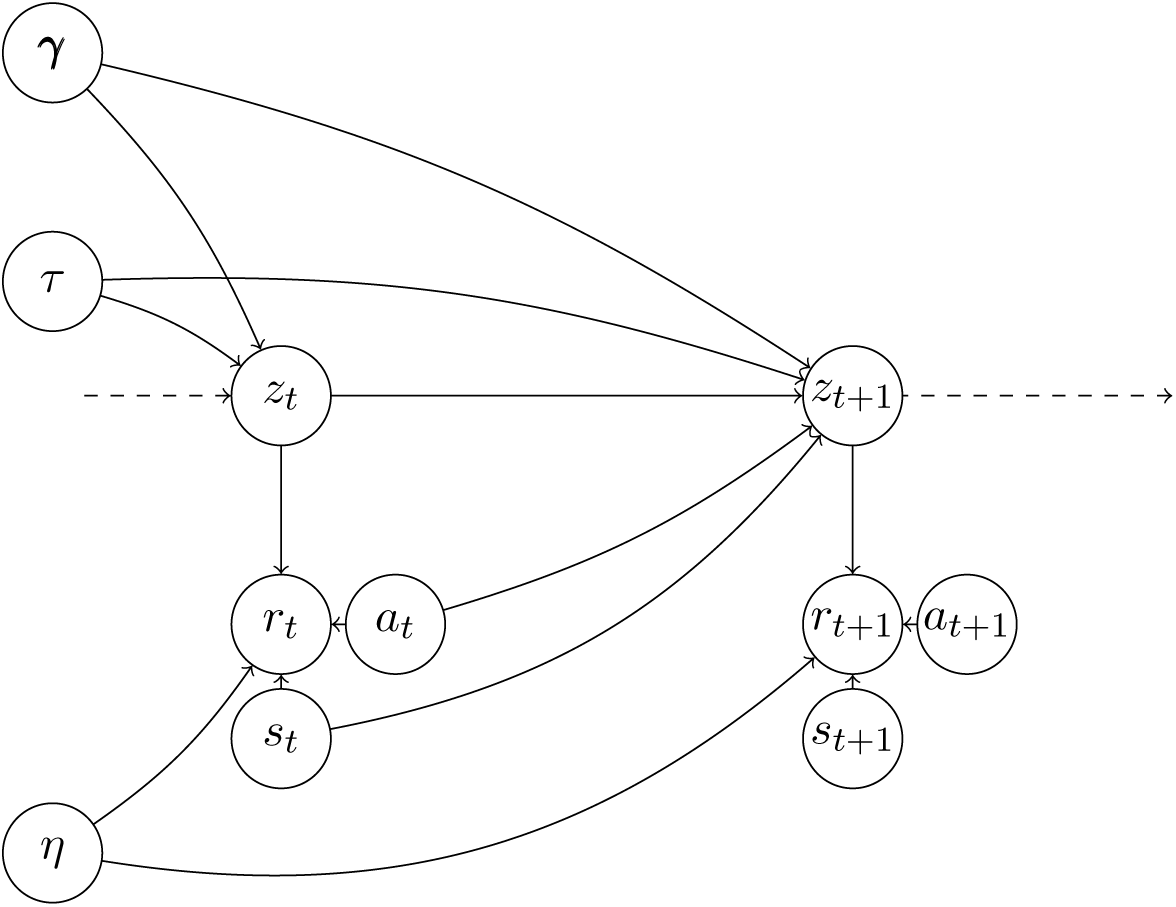
Full generative model of constant-volatility models. This model is exactly the process generating *changing* environments. This generative model assumes that volatility *τ* is constant. *z*_*t*_ represents the current correct combination. *γ* represents the probabilities of combination occurrence whenever the correct combination changes. *η* represents feedback noise. In every trial, observables are stimuli *s*_*t*_, actions *a*_*t*_ and binary feedback *r*_*t*_. See Methods for details.

**Supplementary Figure 5:**
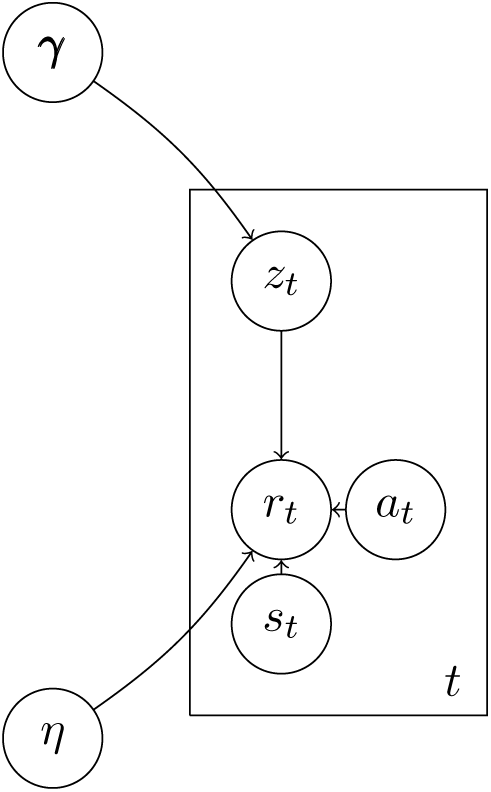
Full generative model of zero-volatility models. This model is exactly the process generating *stable* environments. This generative model assumes that volatility is null and that observations are all equally informative. *z*_*t*_ represents the correct combination. *γ* represents combinations’ probabilities. *η* represents feedback noise. In every trial, observables are stimuli *s*_*t*_, actions *a*_*t*_ and binary feedback *r*_*t*_. See Methods for details.

**Supplementary Figure 6:**
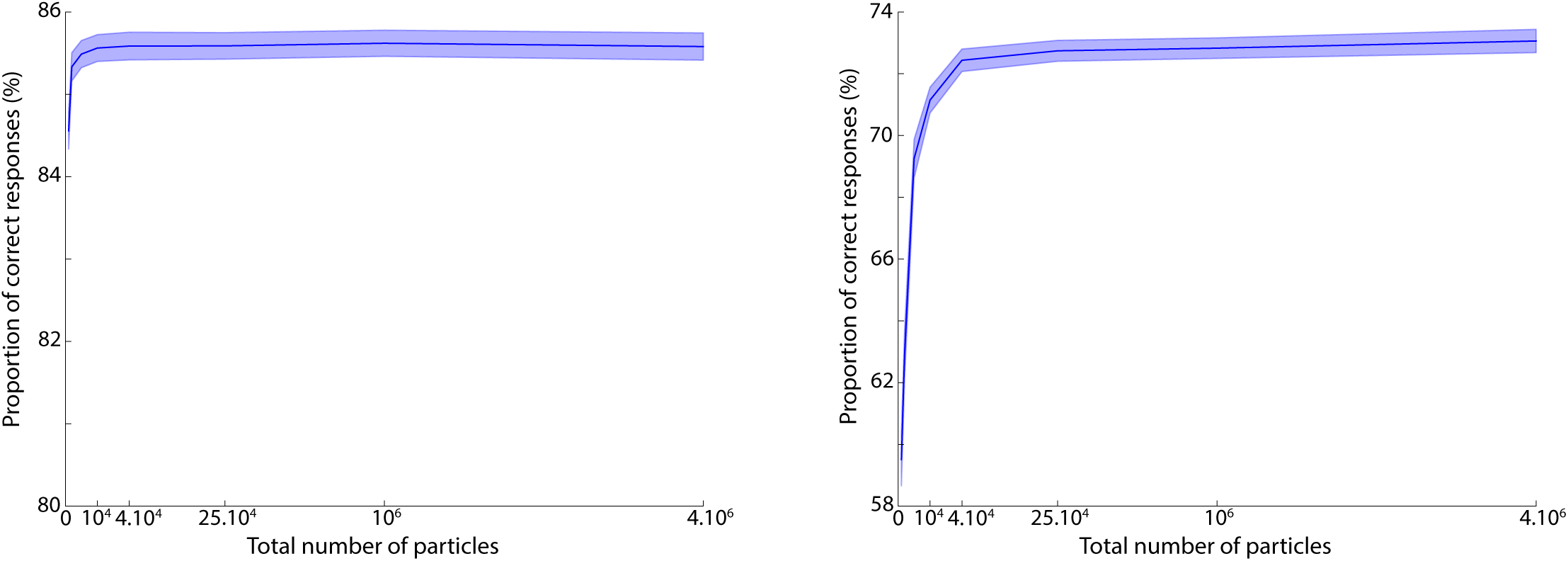
Performance of the forward varying-volatility model in *unstable* environments for different number of particles. The forward varying-volatility model is simulated in *unstable* environments with *N* = *N*_*θ*_ samples for the Sequential Monte Carlo (SMC) procedure in the parameter space, each associated with *N* = *N*_*z*_ samples for the SMC in the latent space leading to a total of *N*_*θ*_ · *N*_*z*_ = *N*^2^ particles. Performance is defined as fraction of times that the agent choses the highest-rewarding action. For a total number of particles greather then 4 · 10^4^, the asymptote performance is reached in both closed and open-ended environments. We thus select 4 · 10^4^ particles for all our experiments. See Methods for more details.

**Supplementary Table 1:** Means and s.e.m (in parentheses) of parameters fitted across subjects for exact and forward volatility models.

| | Softmax coefficient $\beta$ | |
| --- | --- | --- |
| Volatility Model | Closed environments | Open-ended environments |
| Exact Varying Volatility | 1.61 (0.13) | 1.13 (0.021) |
| Exact Constant Volatility | 1.34 (0.12) | 0.94 (0.017) |
| Forward Varying Volatility | 1.63 (0.11) | 1.27 (0.024) |
| Forward Constant Volatility | 1.38 (0.11) | 1.07 (0.02) |

**Supplementary Table 2:** Means and s.e.m (in parentheses) of parameters fitted across subjects for the Weber-imprecision model.

|  | Weber-imprecision Model |  |
| --- | --- | --- |
| Parameter | Closed environments | Open-ended environments |
| Softmax coefficient $\beta$ | 1.77 (0.13) | 1.05 (0.023) |
| Weber Component $\lambda$ | 0.8 (0.1) | 1.22 (0.06) |
| Constant Component $\mu$ | 0.05 (0.01) | 0.05 (0.003) |

**Supplementary Table 3:**
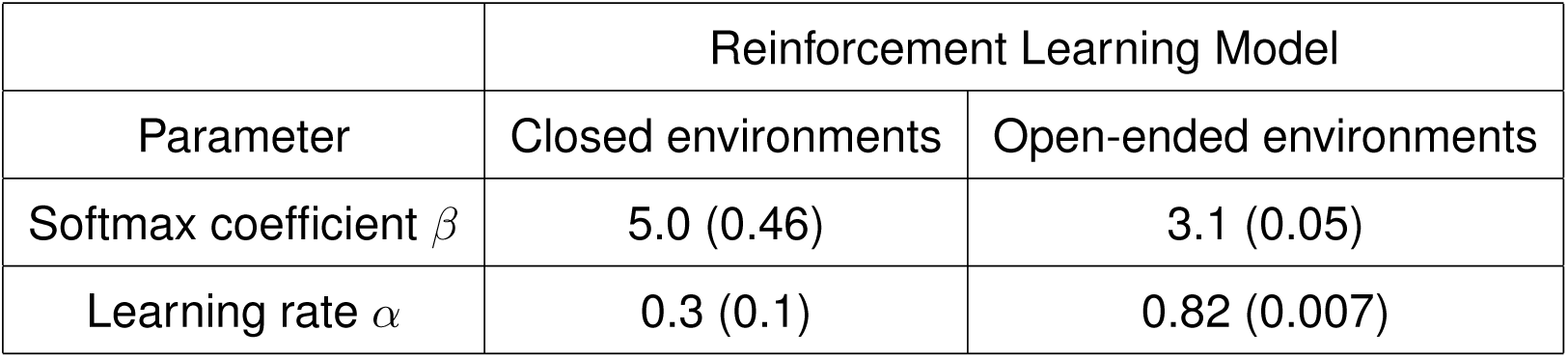
Means and s.e.m (in parentheses) of parameters fitted across subjects for the reinforcement learning model.

|  | Reinforcement Learning Model |  |
| --- | --- | --- |
| Parameter | Closed environments | Open-ended environments |
| Softmax coefficient $\beta$ | 5.0 (0.46) | 3.1 (0.05) |
| Learning rate $\alpha$ | 0.3 (0.1) | 0.82 (0.007) |

### Supplementary Methods

#### 1 Analytical models

The generative processes that define the models have been described in the Main Text and Methods and their graphical representations are among the Supplementary Figures (3-5). Regarding the formal equations, they can be found hereafter.

##### 1.1 Varying-Volatility

Using standard notations, the full varying-volatility analytical model writes as follow:

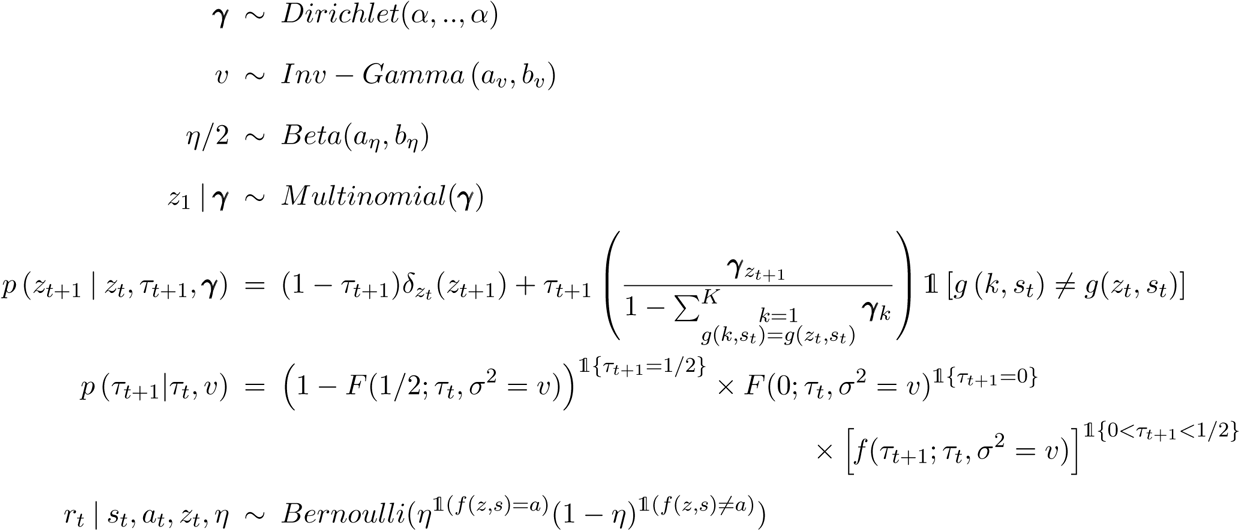

***γ*** represents the distribution over possible stimulus-action combinations after a correct combination change, *τ*_*t*_ is the volatility, *ν* the variance of the volatility random walk and *η* represents feedback noise. *z*_*t*_ is the current correct combination, *r*_*t*_ the feedback, *s*_*t*_ the stimulus and *a*_*t*_ the chosen action. Hyper-parameters’ values correspond to uninformative priors: *α* = 1, *a*_*ν*_ = 3, *b*_*ν*_ = 0.001, *a*_*η*_ = 1 and *b*_*η*_ = 1. For the prior on feedback noise, rç, we thus assume a uniform distribution between 0.5 and 1 (see Main Text).*g* is the stimulus-action mapping function that specifies the correct action *a* for every possible combination *z* and stimulus s: *a* = *g(z, s). F*(.;*m, s)* and *f*(.;*m, s)* are respectively the Gaussian cumulative distribution function and density with mean *m* and standard deviation *s*. 𝟙 denotes the indicator function.

##### 1.2 Constant-Volatility

Using standard notations, the full constant-volatility analytical model writes as follow:

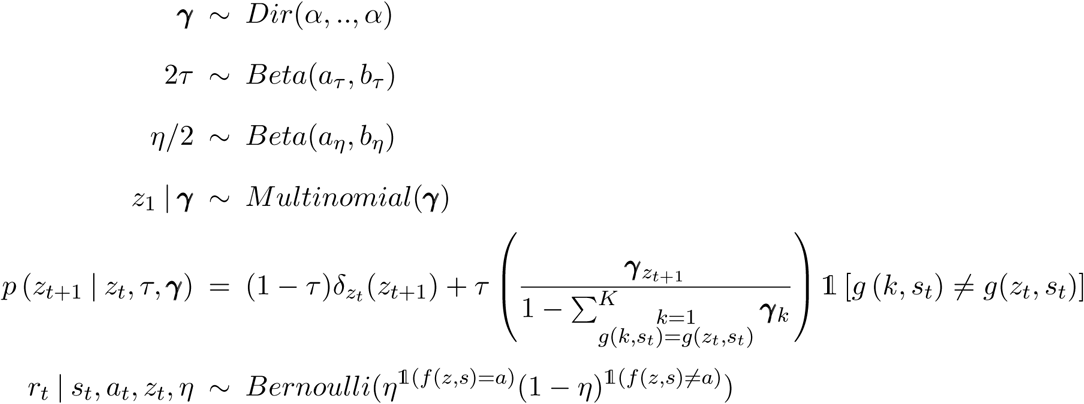

*γ* represents the distribution over possible stimulus-action combinations after a correct combination change, *τ* is the volatility and *η* represents feedback noise. *z*_*t*_ is the current correct combination, *r*_*t*_ the feedback, *s*_*t*_ the stimulus and *a*_*t*_ the chosen action. Hyperparameters’ values correspond to uninformative priors: *α* = 1, *a*_*τ*_ = 1, *b*_*τ*_ = 1, *a*_*η*_ = 1 and *b*_*η*_ = 1. Regarding the priors on volatility *τ* and feedback noise *η*, we assume uniform distributions: the first between 0 and 0.5 and the second between 0.5 and 1 (see Main Text). *g* is the stimulus-action mapping function that specifies the correct action *a* for every possible combination *z* and stimulus s: *a* = *g(z, s). F*(.;*m, s)* and *f*(.;*m, s)* are respectively the Gaussian cumulative distribution function and density with mean *m* and standard deviation *s*.𝟙 denotes the indicator function.

##### 1.3 Zero-Volatility

Using standard notations, the full zero-volatility analytical model writes as follow:

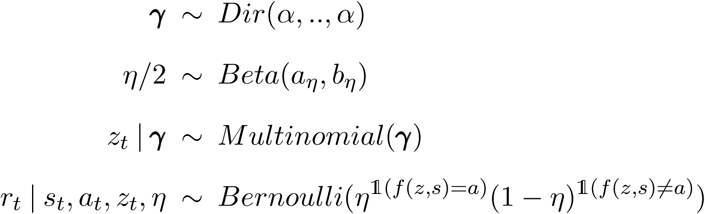

*γ* represents the distribution over possible stimulus-action combinations and *η* represents feedback noise. *z*_*t*_ is the correct combination, *r*_*t*_ the feedback, *s*_*t*_ the stimulus and *a*_*t*_ the chosen action. Observations *r*_*t*_, *s*_*t*_ and *a*_*t*_ are assumed independent and identically distributed. Hyper-parameters’ values correspond to uninformative priors: *α* = 1, *a*_*η*_ = 1 and *b*_*η*_ = 1. For the prior on feedback noise, *η*, we thus assume a uniform distribution between 0.5 and 1 (see Main Text). *g* is the stimulus-action mapping function that specifies the correct action *a* for every possible combination *z* and stimulus s: *a* = *g(z*, s). *F*(.;*m, s*) and *f*(.;*m, s*) are respectively the Gaussian cumulative distribution function and density with mean *m* and standard deviation *s*. 𝟙 denotes the indicator function.

#### 2 Deriving the update quantity from the Weber-imprecision model

The Weber-imprecision model is derived from the zero-volatility model where inference is performed with the same forward procedure as in the forward varying- and constant-volatility model (see Methods). The Weber-imprecision model assumes that, consistent with the Weber law, computational imprecisions in computing posterior belief scale with the distance between posterior beliefs in trial t-1 and t.

Using standard notations, let us write 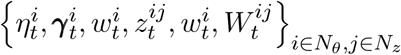 the particle system in trial t. This particle system is composed of *N*_*θ*_ samples 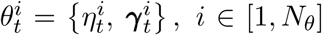 representing the posterior in the parameter space : samples 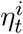 represent the posterior about feedback noise and 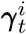 about combinations’ probabilities whenever the correct combination changes. And, for each 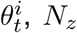 particles 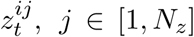 approximate the corresponding latent state posterior beliefs. All particles 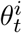 and 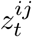 are assigned weights 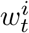 and 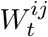 respectively, reflecting theirs likelihoods (see references [1-5] for more details on particle filters). From the particle system, we can compute estimators of posterior beliefs in trial *t* − 1 and *t.* For every *z* ∈ [1,*K*], with *K* the size of the latent state space, these estimators write as follow,

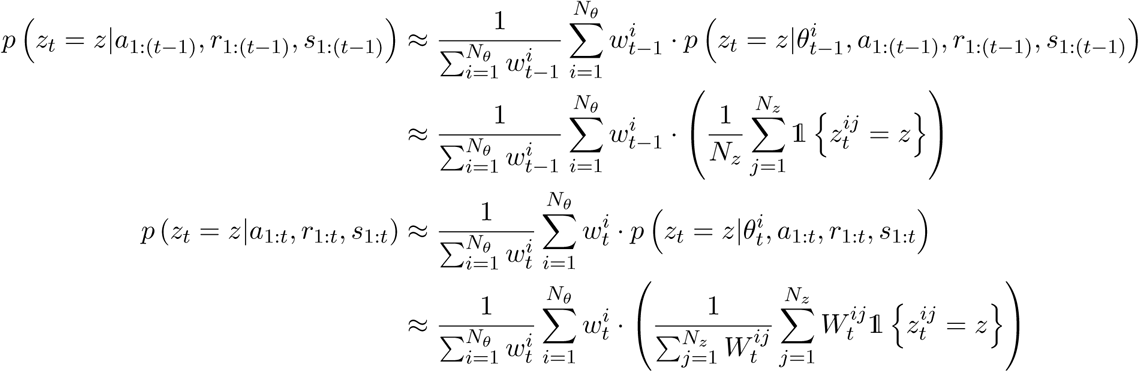

This leads to the distance between posterior beliefs in trial *t* − 1 and *t*:

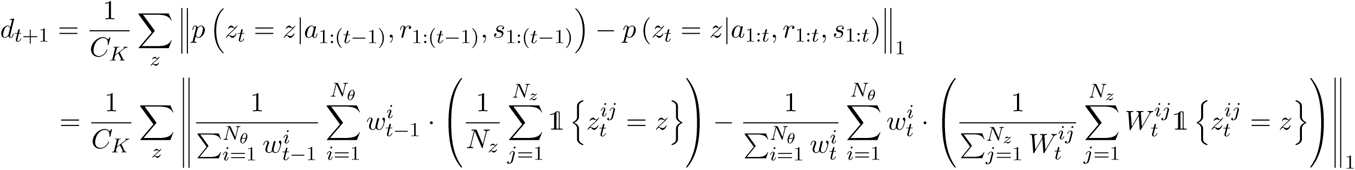

*C*_*K*_ is a normalization factor depending on the number of potential combinations which are orthogonal to the current correct combination (no shared stimulus-action pairs). For closed environments, the two combinations are orthogonal, thus *C*_*K*_ = 2. For open-ended environments, 11 combinations share no stimulus-action pair with the current correct combination such at *C*_*K*_ = 12. This normalization was introduced in order to make parameter *λ* commensurable accross open-ended and closed environments.

#### 3 Deriving the forward resampling step

Instead of performing formal P-MCMC steps, forward models sample naively from empirical distributions based on current particle systems (see Methods). To implement this step, we resample our parameters from 1) Beta distributions – for the volatility *τ* in the forward constant-volatility model and feedback noise *η* in both forward models, 2) Inverse-Gamma distributions – for the variance *ν* of the bounded volatility random walk in forward varying-volatility model, and 3) Dirichlet distributions for *γ*, the probabilities of combination occurence whenever the correct combination changes, in both forward models.

When naive resampling occurs, we thus sample from empirical distributions. To obtain these distributions, we compute their empirical mean and variance from which we derive the distributions’ characteristic parameters. Regarding the derivation of empirical means and variances, obtaining them from the particle system is rather straightforward. For feedback noise for instance, with the notation of the previous paragraph, the empirical mean and variance write as follow:

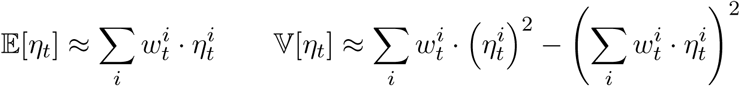

We will now study how to derive the characteristic parameters for our three distributions of interest.

##### 3.1 Beta distribution

Let X follow a beta distribution with parameters *α* and *β*. We can write the mean and variance of *X* as functions of *α* and *β*:

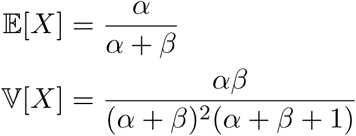

Inverting these equations gives us:

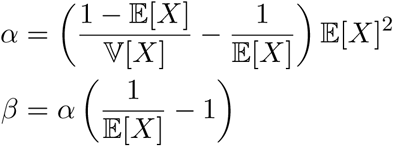

Thus, replacing 𝔼[*X*] and 𝕍[*X*] with their empirical estimates based on the particle system leads to parameters *α* and *β* and thus the Beta proposal.

##### 3.2 Dirichlet distribution

Let *X* = [*X*_1_,…,*X*_*K*_] ∼ *Dirichlet(α*_1_,,…, *α*_*K*_). Again, we can write the mean and variance of each component *X*_*i*_ as functions of the Dirichlet parameters :

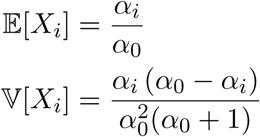

With *α*_0_ = ∑*α*_*i*_. Inverting gives us:

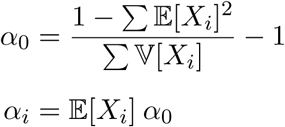

Thus, replacing 𝔼[*X*_*i*_] and 𝕍[*X*_*i*_] with their empirical estimates based on the particle system leads to the Dirichlet proposal.

##### 3.3 Inverse-Gamma

Let *X* ∼ *Inv-Gamma(α, β*). The mean and variance are:

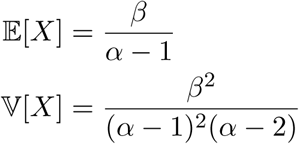

We invert these equations and obtain:

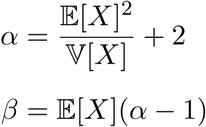

Thus, replacing 𝔼[*X*] and 𝔼[*X*] with their empirical estimates based on the particle system leads to the Inverse-Gamma proposal.

